# A Single-Cell Bioprinting Method to Reconstruct Native Cellular Microenvironments with Subcellular Resolution

**DOI:** 10.1101/2024.02.01.578499

**Authors:** Haylie R. Helms, Kody A. Oyama, Alexander E. Davies, Ellen M. Langer, Luiz E. Bertassoni

## Abstract

Tissue development and function are determined by the spatial organization of individual cells and their interactions. Yet experimental platforms capable of reconstructing cellular organization with single-cell precision for systematic interrogation remain lacking. Here we introduce a single-cell bioprinting platform that enables programmable reconstruction of cellular microenvironments through controlled single-cell placement with down to subcellular spatial control. We demonstrate intercellular spacing down to 1.3 µm, multiplexed deposition of up to eight cell types within a single construct, and reconstruction of biopsy-derived native cellular organization with 99% accuracy for single-cell placement relative to their native tissue coordinates. Using this framework, we show that manipulation of cellular arrangement allows for controlled interrogation of spatial transcriptional programs, cell–cell signaling networks, and migration dynamics in complex tissue microenvironments. This platform provides a generalizable experimental framework for causal interrogation of spatially defined cell–cell interactions at single-cell resolution.

## Main

The fate and function of multicellular systems are strongly influenced by the spatial positioning of individual cells^1–3^. The identity, distance, and topology of neighboring cells determine which interactions occur, which influence cell behavior and shape gene expression^4–8^. Patient samples analyzed via spatial biology tools have increasingly suggested that even micron-scale positional differences in the cell microenvironment can alter transcriptional programs and phenotypic outcomes^9–12^. Although spatial omics technologies now enable high-resolution mapping of these relationships, native tissues remain observational systems in which spatial effects cannot be experimentally controlled or separated from genetic, epigenetic, and systemic influences, limiting dissection of spatial causality.

Existing experimental platforms provide limited control over single-cell placement. Organoids and co-culture models rely on stochastic self-assembly, producing variable cellular neighborhoods across samples. Cell-laden hydrogel bioprinting and lithography-based micropatterning improve spatial control but lack the resolution required to reproducibly specify cell-cell relationships at the single-cell level^2^. Single-cell technologies have been able to manipulate individual cells with precision^13–22^, but have not yet demonstrated precision engineering of heterogeneous multicellular organization at tissue scale and with single cell resolution. These limitations highlight the need for experimental systems in which cellular spatial organization can be specified with single-cell precision, systematically manipulated, and coupled to quantitative molecular and functional measurements to directly test how spatial interactions govern functional outcomes. Such capabilities would establish a new class of experimental platforms that enable tissue-derived spatial hypotheses to be reconstructed and interrogated under controlled conditions.

Here we introduce a single-cell bioprinting method that enables programable placement of individual cells with subcellular spatial precision. This capability allows multicellular systems to be constructed with defined cell types, spatial relationships, and neighborhood topologies at native tissue scale. Using cellular coordinates derived from patient tissue, the platform reconstructs native cellular microenvironments with over 99% spatial accuracy while remaining compatible with diverse cell types, substrates, and analytical workflows. By integrating quantitative imaging and spatial transcriptomics with engineered tissues, variables such as cellular composition and spatial organization can be systematically manipulated and linked to cellular behavior and molecular readouts (**Fig. 1A**). Using this framework, we show that controlled perturbation of microscale cellular organization is sufficient to produce reproducible changes in migration trajectories, tissue architecture, transcriptional programs, and inferred cell–cell signaling networks that were not observed against benchmarked samples prepared using standard methodologies. Together, this approach establishes an experimental platform for controlled assembly of multicellular systems, enabling systematic interrogation of how intrinsic and microenvironmental variables govern tissue behavior at the single-cell level.

**Figure 1.**
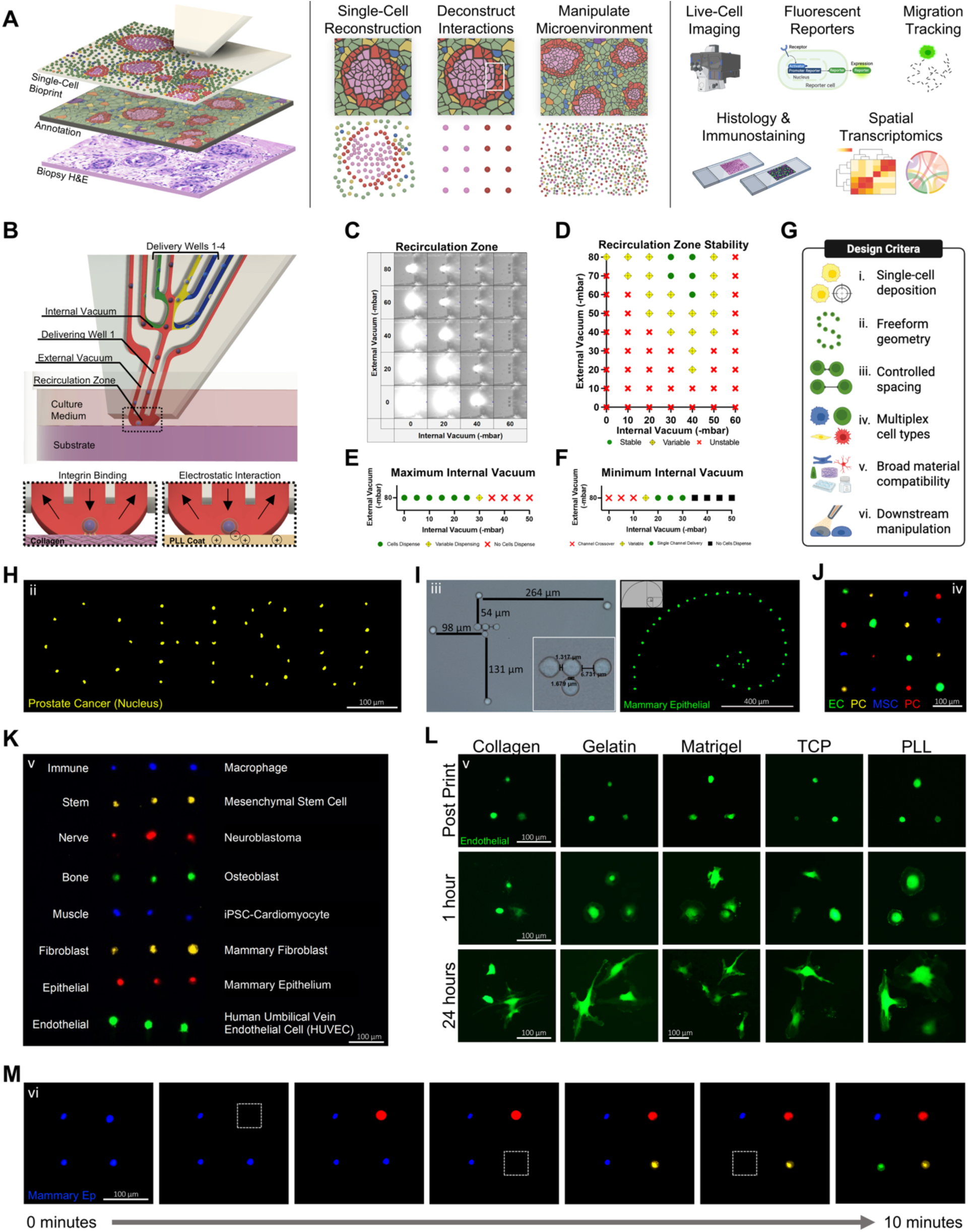
Development and validation of controlled single-cell bioprinting. **A)** Conceptual overview of the single-cell bioprinting workflow for reconstruction and perturbation of cellular microenvironments followed by downstream imaging and molecular analysis. H&E obtained from 10x Genomics public human breast dataset^47^; right created with BioRender.com. **B)** Microfluidic printhead schematic showing internal fluidic channels, vacuum interfaces, cell delivery and cell attachment at the substrate surface. **C)** Visualization of the recirculation zone using fluorescein flowing through transparent medium at 80 mbar delivery pressure. **D)** Recirculation zone stability across different printheads at 80 mbar delivery pressure (*n* = 3 tests per printhead). **E)** Maximum internal vacuum permitting cell dispensing at 80 mbar delivery pressure (*n* = 9 tests across 3 printheads). **F)** Minimum internal vacuum preventing unintended dispensing from neighboring wells at 80 mbar delivery pressure (*n* = 9 tests across 3 printheads). **G)** Design criteria guiding development of the single-cell bioprinting platform. Created with BioRender.com **H–M)** Experimental validation of the design criteria shown in (G), corresponding to criteria i–vi indicated within each panel. **H)** Deterministic single-cell deposition. Each target location contains one deposited cell without doublets or missing positions. Prostate cancer cells labeled with a fluorescent nuclear stain were imaged immediately after printing before cell spreading. Scale bar, 100 µm. **I)** Controlled intercellular spacing demonstrated using a Fibonacci spiral of mammary epithelial cells. Image acquired immediately after printing. Scale bar, 400 µm. **J)** Multiplexed deposition of four distinct cell populations within a single print run. Image acquired immediately after printing. EC, endothelial cell; PC, prostate cancer; MSC, mesenchymal stem cell. 100 µm. **K)** Compatibility across diverse cell types. Eight representative human cell types patterned within a single bioprint and fluorescently labeled prior to printing for identification. Image acquired immediately after printing. Scale bar, 100 µm. **L)** Compatibility across substrates. GFP+ human umbilical vein endothelial cells bioprinted onto collagen, gelatin, Matrigel, tissue culture plastic (TCP) and poly-L-lysine–coated plastic. Scale bars, 100 µm. **M)** Post-fabrication manipulation of individual cells using the recirculation zone. A single mammary epithelial cell was selectively detached using TrypLE and replaced with a newly deposited cell at the same coordinate. Scale bar, 100 µm.

## Results

### Engineering parameters enabling single-cell bioprinting

Experimentally dissecting spatial interactions within tissues requires controlled and reproducible placement of individual cells at native tissue scale and single-cell spatial precision, a capability not demonstrated by existing approaches (**Table S1**). We therefore defined the following technical criteria: i) deposition of exactly one cell per target location, ii) flexible patterning of spatial geometries rather than fixed grid patterns, iii) controllable intercellular spacing below the diameter of a detached cell (∼18 µm) to achieve cellular densities comparable to native tissues, and iv) multiplexed deposition of multiple cell types within a single print run for heterogeneous tissue fabrication. To ensure broad experimental utility, the platform was additionally required to be compatible with diverse cell types and biomaterials and support downstream experimental workflows including live-cell manipulation (i.e., live-seq^23^, single-cell dosing^24^, patch clamp), imaging, and molecular profiling.

To identify a platform capable of meeting these requirements, we benchmarked several single-cell manipulation technologies described in the literature and available in our laboratory, including laser-assisted forward transfer bioprinting (NGB-R, Poietis), inkjet single-cell dispensing (D100, Hewlett-Packard), robotic single-cell picking (CellCelector, Sartorius), and open-volume microfluidic cell dispensing (Biopixlar, Fluicell) (**Table S1**). From the available systems, the open-volume microfluidic cell dispenser most closely satisfied the multiplexing and flexible fabrication requirements. We therefore sought to identify a set of engineering and hardware parameters that would allow controlled and reproducible transport of heterogeneous cell singlets for patterning of virtually unlimited spatial configurations on substrates ranging from hydrogels to tissue culture plastic.

The flow of the microfluidic dispenser is controlled by four independent parameters (delivery pressure, non-delivery pressure, internal vacuum, and external vacuum), accounting for more than two billion possible parameter combinations that remained to be defined for heterogeneous single-cell bioprinting (**Fig. 1B and S1**). We systematically screened these parameter combinations to identify conditions that produced a stable recirculation zone for single-cell transport and deposition. Detailed descriptions of device fluidics and parameter optimization are provided in **Supplementary Information File 2**. Initial experiments were performed using cell culture medium alone to identify parameter regimes supporting stable fluid interfaces (**Fig. 1C–D and S2; Video 1**). Cells were subsequently introduced to determine which conditions supported consistent single-cell delivery (**Figs. 1E and S3**) while preventing unintended cross-dispensing between reservoirs (**Figs. 1F and S3**).

To maximize positional accuracy following dispensing, we next optimized substrate attachment conditions to prevent lateral cell drift after deposition. Hardware modifications included maintaining the culture chamber at 37 °C to support rapid cell–substrate adhesion, increasing printhead angulation to promote immediate contact with the substrate, and implementing custom software that overlays the bioprint design onto the live microscope feed to guide printhead positioning relative to the intended pattern. Using these modifications, primary human umbilical vein endothelial cells (HUVECs) were bioprinted onto collagen substrates and evaluated based on the binding success, ability to deposit a single cell vs doublet(s), and bioprinting speed (**Fig. S4**).

Across these experiments we identified an optimal operating regime of 80 mbar delivery pressure, 0 mbar non-delivery pressure, −25 mbar internal vacuum, and −40 mbar external vacuum, combined with a 70° printhead angle and chamber temperature control. Under these conditions, the platform reproducibly deposited individual cells with precise spatial control, establishing a fluidic regime for controlled and reproducible single-cell bioprinting.

### Single-cell bioprinting enables micron-scale spatial control, multiplexed patterning, and broad compatibility

We next tested whether the platform could generate spatially defined architectures while satisfying the design criteria outlined above (**Fig. 1G**). Because reconstruction of complex tissues requires simultaneous control of cell position, spacing, and heterogeneity, we first validated these capabilities individually before integrating them to fabricate complex multicellular tissues. As an initial validation of positional accuracy and geometric flexibility, individual cells were bioprinted to write the acronym of our university (“OHSU”) (**Fig. 1H**). Each target position contained a single deposited cell with no missing locations or doublets. We then examined the spatial resolution achievable for intercellular positioning by bioprinting the Fibonacci spiral sequence (**Fig. 1I**). At the smallest positions, cells were deposited with edge-to-edge intercellular spacing of 1.3 µm.

Because native tissues contain multiple interacting cell populations, we next assessed multiplexed printing capability. Unlike other high precision cell patterning strategies, which can only support one cell type at a time, this platform enables simultaneous deposition of up to four different cell populations without having to swap the loaded material. Endothelial cells, mesenchymal stem cells, and two distinct prostate cancer cell populations were loaded into the printhead and bioprinted into an array of alternating cell populations (**Fig. 1J**). This confirmed that the optimized fluidic regime reliably delivered the correct cell population without cross-dispensing between reservoirs, enabling heterogeneous multicellular architectures to be fabricated within a single print run.

Reconstruction of tissue microenvironments further requires compatibility across diverse cell types and biomaterials. To assess biological compatibility, we bioprinted eight representative cell lineages (immune, stem, neural, bone, muscle, fibroblast, epithelial, and endothelial) onto a collagen substrate in a single sample using two printheads (**Fig. 1K**). The established bioprinting conditions supported reliable deposition across all tested cell types despite differences in cell size, morphology and aggregation behavior that can perturb fluid flow, demonstrating robustness of the printing regime. We then evaluated compatibility with commonly used biological substrates. HUVECs were bioprinted onto collagen, gelatin, Matrigel, tissue-culture plastic and poly-L-lysine–coated plastic (**Fig. 1L**). Cells adhered to all substrates and exhibited spreading and migration within one hour of deposition, indicating preservation of cell health following printing. Occasional cell division was also observed within this period, suggesting that the bioprinting process did not induce sufficient stress to disrupt cell-cycle progression. Imaging at 24 hours confirmed expected morphology and migration across groups.

Finally, to demonstrate compatibility with downstream experimental workflows, the fabrication strategy was designed to pattern cells without requiring cell encapsulation, preserving access to individual cells for post-fabrication manipulation or sampling. Using the confined recirculation zone generated by the printhead (**Video 1**), we selectively detached a single printed cell by locally delivering TrypLE while leaving neighboring cells undisturbed^24^ (**Fig. 1M**). The detached cell was aspirated through the external vacuum channel and replaced with a new cell deposited at the same location. This capability enables targeted post-fabrication manipulation of individual cells within engineered multicellular systems. In addition, cells can be encapsulated in hydrogel following printing, allowing three-dimensional constructs to be fabricated sequentially, one layer at a time (**Fig. S5**).

### Reconstruction of patient tissue cellular organization with subcellular resolution

Having established controlled single-cell placement, spatial resolution, and multiplexing capability, we next asked whether these capabilities could be combined to reconstruct spatial organization derived from patient tissues. A region of interest from a previously annotated human breast cancer biopsy^25^ was selected, containing malignant mammary epithelial cells with surrounding stromal compartment (**Fig. 2A**). Cellular coordinates were extracted from the annotated dataset and converted into a bioprinting map for placement of five distinct cell populations: mammary epithelial cells, triple negative breast cancer cells, mammary fibroblasts, mesenchymal stem cells, and macrophages. Across four bioprints, the mean absolute distance between intended target position, defined by the bioprint map, and actual bioprinted cell position was 3.14 µm, with 99.62% of points within one cell diameter (**Figs. 2B-E and S6A**). Compared to published methods that report positional accuracy within 10 µm of intended target^16^, but rely on substantially larger spacing (i.e., 400 µm^16^) to achieve reproducible placement, our approach enables high-fidelity reconstruction of native cellular microenvironments while preserving both cellular identity and spatial organization. To the best of our knowledge, these findings establish the first bioprinting methodology capable of reconstructing native cellular microenvironments with near exact spatial accuracy.

**Figure 2.**
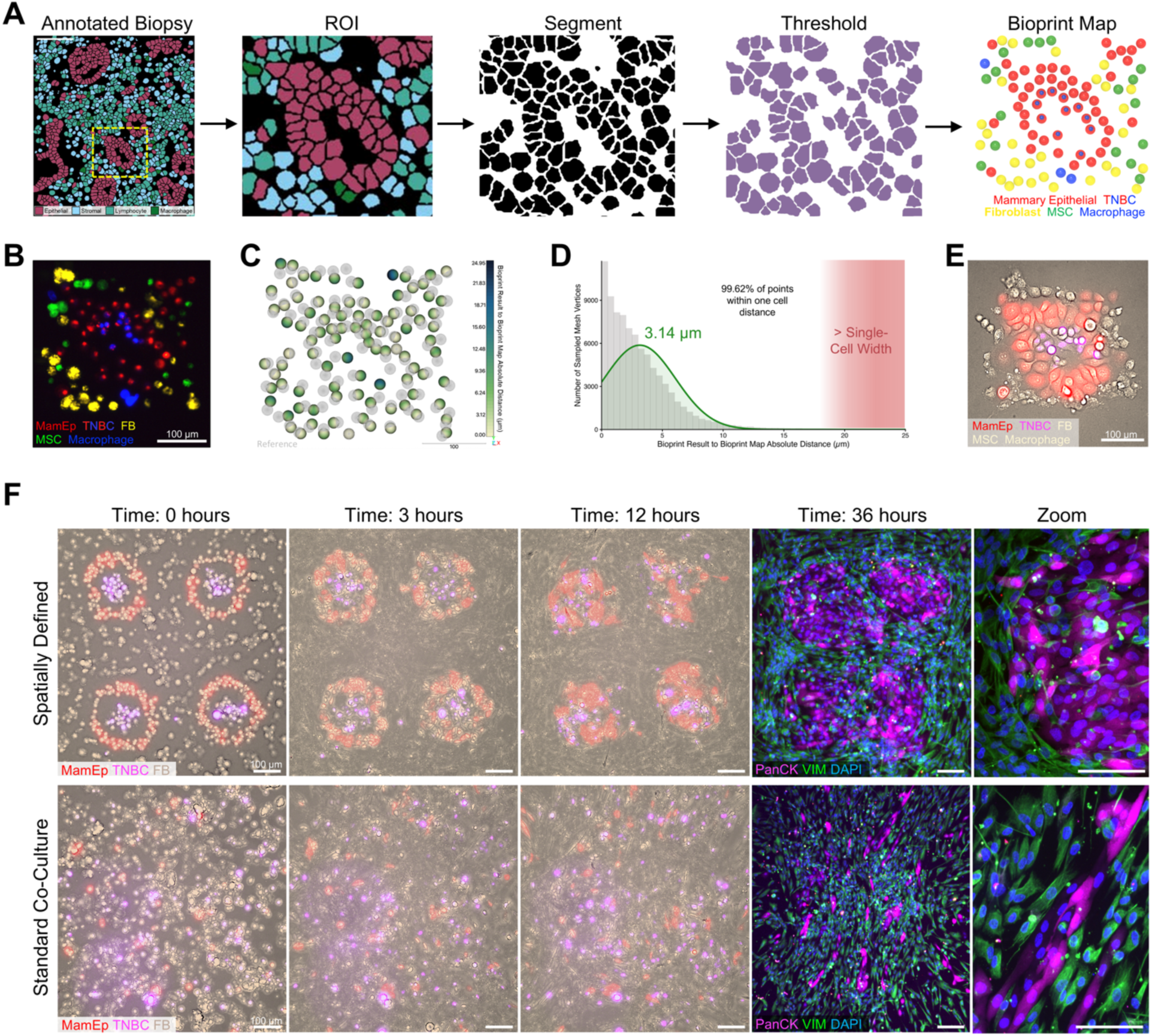
Reconstruction of biopsy-derived cellular organization and effects of spatial patterning on tissue assembly. **A)** Workflow for generating a single-cell bioprint map from an annotated breast cancer biopsy. Image adapted with permission^25^ 2020 Springer Nature. Scale Bar, 100 µm. **B)** Bioprinted reconstruction of the biopsy-derived cellular organization. Cell populations were fluorescently labeled prior to printing: mammary epithelial (red), triple negative breast cancer (red + blue), fibroblasts (yellow), mesenchymal stem/stromal cells (green) and macrophages (blue). Scale bar, 100 µm. **C)** Fidelity analysis comparing printed cell positions to the reference bioprint map (gray). Colors indicate spatial deviation (µm) between printed cells and target coordinates. Scale bar, 100 µm. **D)** Distribution of spatial deviations between printed cells and the reference bioprint map across independent samples. Absolute distances were computed between reconstructed cell positions and target coordinates. ∼18 µm indicates one detached-cell diameter. Mean spatial deviation 3.14 ± 2.93 µm. (*n* = 4 bioprints). **E)** Reconstruction of the same bioprint map using genetically labeled cells without cell-tracker dyes (EF1α–mammary epithelial cell, red; H2B–triple negative breast cancer, pink). Scale bar, 100 µm. **F)** Comparison of spatially defined tumor microenvironment and randomly deposited co-culture bioprints containing the same cell ratios (mammary epithelial, triple negative breast cancer, and mammary fibroblasts). Left: representative live-cell imaging timepoints following printing. Right: endpoint immunofluorescence staining for pan-cytokeratin (PanCK), vimentin (VIM) and DAPI. Scale bars, 100 µm.

Live-cell imaging of the reconstructed tumor microenvironment confirmed preserved cell viability, with cells rapidly adopting expected morphologies, migrating, and proliferating over time (**Video 2; Fig. S6B**). Mammary epithelial cells exhibited characteristic morphology and cell–cell contacts resembling the epithelial structures observed in the original annotated native tissue (**Fig. 2E**). Over time, cells began migrating outward into unoccupied regions of the substrate, spreading in the absence of physical constraints that normally limit tissue expansion in vivo. We therefore reasoned that increasing local cellular density would promote architectural stability. To test this, we bioprinted an idealized tissue configuration consisting of an epithelial ring surrounding triple negative breast cancer cells, encased within a dense stromal fibroblast compartment (**Fig. S6C-D**). Immunostaining at 72 hours confirmed the persistence of epithelial ring-like structures surrounded by stromal cells (**Fig. S6D**), demonstrating that densely patterned architectures can support prolonged maintenance of prescribed spatial organization.

### Spatial organization drives divergent architecture, transcriptional programs, and signaling networks

Standard in vitro models rely on the capacity of cells to self-organize into tissue-like structures, raising the question of whether explicit single-cell spatial control is necessary. To address this, we performed a direct comparison between bioprints with defined cellular spatial organization and benchmarked it against standard cell co-culture experiments in which the same cell ratios were used without imposing spatial structure (**Figs. 2F; Videos 3-4**). To ensure that the effect of bioprinting was accounted for in both groups, we utilized the same cell dispenser to either define cellular spatial organization or to generate random cellular organization, as typically occurring in conventional co-culture experiments. Despite the same starting cell ratios, the two conditions diverged markedly in cellular morphology, organization, and local density. Spatially defined bioprints preserved epithelial architecture, with fibroblasts organizing circumferentially around epithelial nest-like structures. In contrast, co-cultures failed to self-assemble into hallmark architectural features of native tissues. Fibroblast elongation appeared to bias the migration of neighboring epithelial cells in co-culture constructs, disrupting the characteristic epithelial cobblestone morphology maintained in spatially defined bioprints.

To further investigate the effect of spatial patterning on cell state and transcriptional signature, we performed single-cell spatial transcriptomic analysis (Visium HD) on spatially defined tumor microenvironments and conventional co-cultures generated from the same starting cell populations (**Figs. 3 and S7**). Known cell identities were assigned using endpoint fluorescence imaging in which each cell type expressed a unique fluorescent reporter, enabling direct correspondence between spatial transcriptomic barcodes and starting cell identities (**Figs. 3A and S7A–B**). Conventional computational annotation approaches deviated substantially from these fluorescence-defined ground truth labels (**Fig. S7C**), therefore downstream analyses were performed using the fluorescence-based annotations. Although both conditions were initiated with the same cell type ratios, endpoint population analysis revealed divergence in cellular composition in co-culture constructs, characterized by increased representation of the cancer population and reduced representation of mammary epithelial cells (**Fig. 3B**). This highlights a limitation of conventional co-culture models, which lack reproducible control over cellular neighborhoods, and suggests that random organization may preferentially support cancer expansion compared to spatially defined constructs in which healthy epithelial cells surrounded the cancer cells. Expression of canonical marker genes derived from internal single-population controls mapped as expected across the spatial transcriptomic dataset, further supporting the manual cell-type annotations (**Figs. 3C and S7D–E**).

**Figure 3.**
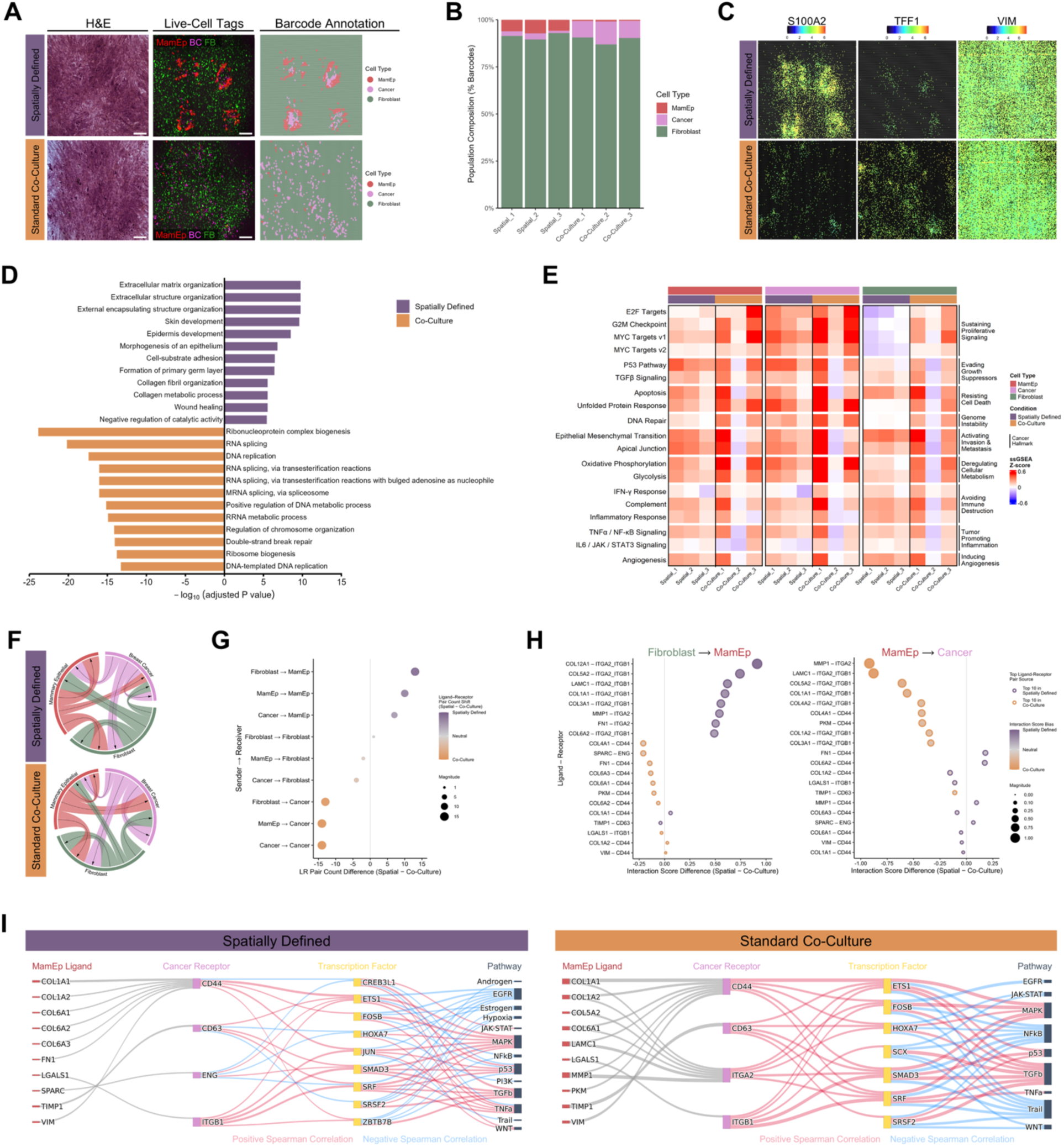
Spatial organization alters gene expression programs and inferred cell–cell signaling networks. **A)** Representative spatially defined and randomly deposited co-culture bioprints containing mammary epithelial cells (EF1α–tdTomato9+), breast cancer cells (H2B–iRFP+) and mammary fibroblasts (H2B–mVenus+). Hematoxylin and eosin staining (H&E), live fluorescence images of labeled cells prior to fixation, and barcode-level cell-type annotation used for spatial transcriptomic analysis. (*n* = 3 bioprints per group). **B)** Proportion of spatial transcriptomic barcodes assigned to each cell type in spatially defined and standard co-culture bioprints across replicates. **C)** Spatial expression patterns of selected marker genes derived from single-population controls shown for spatially defined and co-culture bioprints. **D)** Gene Ontology Biological Process (GO-BP) enrichment analysis of genes upregulated in spatially defined and co-culture bioprints, displayed as mirrored −log10(adjusted P value) following Benjamini–Hochberg correction. **E)** Single-spot gene set enrichment analysis (ssGSEA) Z-scores for selected MSigDB Hallmark cancer pathways across cell types and spatial conditions. **F)** Inferred ligand–receptor communication networks between cell populations. Edge weights represent the number of ligand–receptor pairs connecting each sender–receiver pair. **G)** Changes in directed signaling interactions between spatially defined and co-culture constructs, quantified as the difference in ligand–receptor pair counts between cell-type pairs. **H)** Differentially ranked ligand–receptor interactions mediating communication between selected sender–receiver cell populations in spatially defined and co-culture bioprints. **I)** Integration of ligand–receptor interactions with inferred transcription factor and pathway activities in receiver cells, illustrating downstream signaling architectures associated with spatial organization. Additional details provided in the source data.

We first asked whether spatial organization reshapes transcriptional programs at the tissue scale. We performed Gene Ontology Biological Process (GO-BP) enrichment analysis on genes differentially expressed between spatially defined and co-culture (**Fig. 3D**). Spatially defined constructs were enriched for programs associated with tissue organization and homeostasis, including extracellular matrix organization, epithelial morphogenesis, and cell–substrate adhesion. In contrast, co-culture constructs were dominated by proliferation- and RNA-processing-associated signatures, including DNA replication, RNA splicing, and mitotic cell-cycle progression.

Because these differences emerged despite the same starting cell populations differing only in spatial neighborhood organization, we then examined whether pathway activity within individual cell populations differed according to their spatial context. Single spot gene set enrichment analysis (ssGSEA) of cancer hallmark^26^ pathways revealed environment dependent shifts across all three cell populations (**Figs. 3E and S7F-I**), with co-cultures displaying greater variability across replicates. In spatially defined constructs, mammary epithelial cells were positioned to surround the cancer compartment and exhibited higher mean apical junction pathway activity compared to co-culture (**Figs. S7F-G**). Similarly, breast cancer cells surrounded by mammary epithelial cells in spatially defined constructs expressed higher mean P53 pathway activity relative to co-culture (**Figs. S7F,H**). In contrast, co-culture constructs showed consistent enrichment of proliferative pathways across all cell types, including E2F targets, G2M checkpoint, and MYC targets, with the strongest and most consistent increases observed in fibroblasts (**Figs. S7F,I**). Comparison to internal single-population controls confirmed that these pathway shifts reflect environmental influences rather than cell-type-intrinsic transcriptional programs (**Figs. S7F-I**).

Given the pronounced transcriptional differences between spatially defined and co-culture constructs, we next asked whether altered intercellular communication networks could explain these differences. Ligand–receptor inference using LIANA^27^ provided evidence that spatial organization significantly reshaped intercellular communication networks (**Figs. 3F–I**). Across all inferred cell–cell interactions, spatially defined constructs exhibited enrichment of signaling toward mammary epithelial cells, whereas co-culture constructs showed increased cancer-directed signaling interactions (**Figs. 3F-G**). Examination of individual predicted sender–receiver relationships further showed that spatial organization altered the specific ligand–receptor pairs preferentially mediating intercellular communication (**Fig. 3H**).

To contextualize how these ligand–receptor differences propagate downstream, we integrated ligand–receptor predictions with transcription factor^28,29^ and pathway activity^30^ inference (**Fig. 3I**). In spatially defined constructs, signaling from mammary epithelial cells to breast cancer cells was associated with positive coupling to WNT and JAK–STAT pathway activity within cancer cells. In contrast, co-culture organization exhibited stronger negative coupling to TRAIL (TNF–related apoptosis-inducing ligand) and positive coupling to TGFβ pathway activity. Together, these analyses provide evidence that controlled microscale spatial organization is sufficient to reshape intercellular communication networks and their downstream transcriptional programs.

### Controlled perturbation of cellular arrangement and composition reveals spatially dependent migration behaviors

While engineered tissues capture spatial interactions within multicellular environments where many variables act simultaneously, simplified multicellular circuits provide a complementary approach for isolating specific spatial relationships and directly examining their effects on cell behavior. To determine whether spatial organization alone is sufficient to influence cell behavior, triple-negative breast cancer and endothelial cells were patterned into arrays with fixed intercellular spacing and composition, varying only the relative configuration into segregated, clustered, or mixed arrangements (**Fig. 4A**). This design was motivated by clinical observations that dispersed versus clustered tumor architectures associate with differences in cancer cell plasticity and patient survival.^31,32^ Live-cell imaging revealed highly dynamic endothelial cell behavior, characterized by frequent interactions with cancer cells and exploratory protrusions consistent with environment sensing (**Fig. 4B; Videos 5-7**). Aggregated migration data across replicates revealed distinct and reproducible spatially dependent patterns. While total distance traveled did not differ significantly across arrangements within a given cell type (**Fig. 4C**), spatial organization strongly influenced migration behavior (**Figs. 4D-H and S8A-C**). For example, in segregated configurations, endothelial cells exhibited persistent, directed migration toward the cancer compartment, whereas in mixed configurations, endothelial cell migration was largely nondirectional, reflected by minimal net center-of-mass displacement.

**Figure 4.**
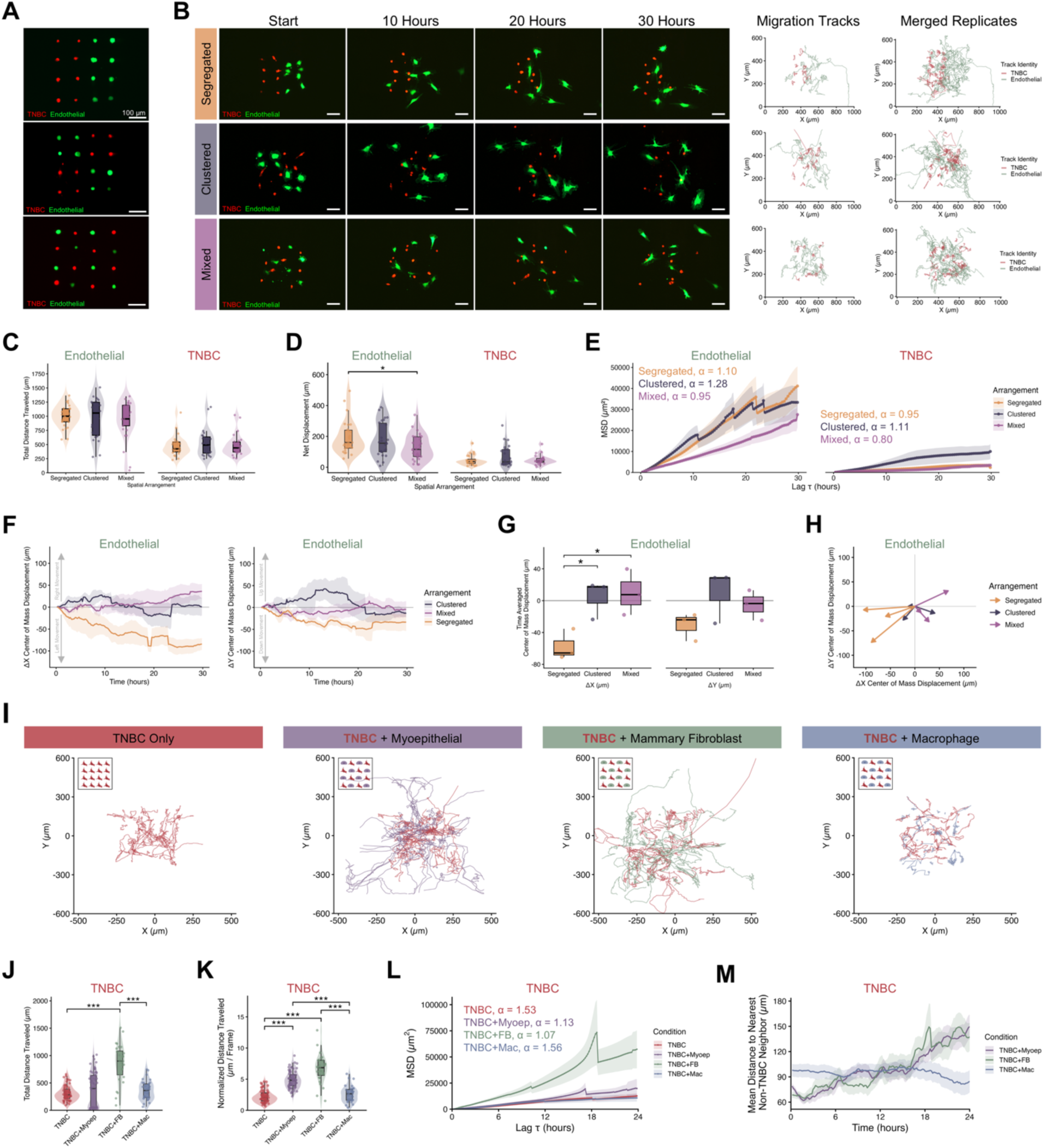
Spatial arrangement and cellular composition influence single-cell migration behavior. **A)** Controlled patterning of individual cells in segregated, clustered or mixed configurations. Triple-negative breast cancer cells (TNBC, CellTracker-labeled) and endothelial cells were imaged immediately after bioprinting. Scale bars, 100 µm. **B–H)** Effect of spatial arrangement on migration behavior in TNBC-endothelial cell interaction arrays. Data are from n = 3 independent bioprinted arrays per spatial configuration. Each point in C and D represents one tracked cell trajectory. Endothelial cells, n = 27, 26 and 29 trajectories; TNBC cells, n = 24, 24 and 26 trajectories for segregated, clustered and mixed arrangements, respectively. **B)** Representative time-lapse fluorescence images with corresponding single-cell migration trajectories. Scale bars, 100 µm. **C,D)** Total distance traveled (**C**) and net displacement (**D**) of individual endothelial and TNBC cells across spatial arrangements. **E)** Time-averaged mean squared displacement (TASD) for each spatial arrangement, plotted as mean ± s.e.m. across tracked cell trajectories. The anomalous diffusion exponent (α) indicates migration mode, with α < 1 indicating subdiffusive migration, α = 1 indicating random migration and α > 1 indicating persistent migration. **F)** Endothelial center-of-mass displacement over time along the X and Y axes, plotted as mean ± s.e.m. across independent bioprinted arrays. **G)** Time-averaged endothelial center-of-mass displacement across spatial configurations. **H)** Center-of-mass displacement vectors for endothelial populations in each array, indicating the direction and magnitude of collective displacement in the X-Y plane. **I–M)** Effect of cellular composition on TNBC migration in mixed-configuration arrays containing myoepithelial cells, fibroblasts, macrophages or TNBC controls at 100 µm spacing. Data are from n = 4 independent arrays per group, except TNBC + fibroblast, where n = 3 arrays. Each point in J and K represents one tracked TNBC cell trajectory, with n = 64, 68, 28 and 34 trajectories for TNBC, TNBC + myoepithelial, TNBC + fibroblast and TNBC + macrophage conditions, respectively. **I)** Combined migration trajectories for TNBC cells in each microenvironment condition. **J,K)** Total distance traveled (**J**) and distance traveled normalized by observation time (**K**) for TNBC cells across microenvironment composition conditions. **L)** TASD of TNBC cells for each microenvironment condition, plotted as mean ± s.e.m. across tracked cell trajectories. **M)** Mean distance from each TNBC cell to its nearest non-TNBC neighbor over time, plotted as mean ± s.e.m. For C, D and G, statistical significance was assessed using one-way ANOVA followed by two-sided Tukey post hoc tests. For J and K, global differences were assessed using Welch’s one-way ANOVA followed by two-sided pairwise Welch t-tests with Benjamini-Hochberg correction. *P < 0.05, **P < 0.01, ***P < 0.001. Exact P values and statistical details are provided in the source data.

Finally, we examined how altering microenvironment cellular composition influences cancer cell migration. Triple negative breast cancer cells were patterned in the same array design either alone or in mixed configuration with myoepithelial cells, mammary fibroblasts, or macrophages and live-cell imaged (**Figs. 4I–M and S8D-G; Videos 8-11**). Changes in microenvironment cellular composition produced distinct cancer cell migration behaviors, reflected by differences in total distance traveled, mean squared displacement, and mean distance to nearest non-cancer neighbor. Taken together, these results demonstrate that precision-engineered microenvironments defined at the level of individual cells provide a controllable framework for systematically manipulating microenvironmental variables to interrogate their influence on cell behavior.

## Discussion

Spatial profiling technologies have greatly enhanced our ability to map cellular organization in tissues, yet platforms that allow systematic manipulation of these spatial relationships with single-cell precision remain limited. Here we present a single-cell bioprinting framework for programmable assembly of multicellular microenvironments, achieving micron-scale control over the identity and position of individual cells in a complex multicellular microenvironment. While manipulation and dispensing of single cells in vitro has been a reality in the field, we argue that this approach for the first time allows one to recreate heterogeneous tissue structures with single cell resolution and 99% accuracy in the laboratory, which represents a leap in performance for bioengineered tissues in vitro. We argue that deterministic assembly of multicellular systems using this method can expand the experimental design space for probing how individual cell behaviors collectively shape tissue organization and function. More broadly, programmable spatial assembly provides a framework for testing hypotheses generated by spatial omics studies and identifying the spatial rules governing cell–cell communication and cell state transitions.

A key advance of this work is the integration of engineered tissues fabricated at single-cell resolution with high-definition spatial transcriptomic analysis. To enable this workflow, we adapted a whole-transcriptome spatial transcriptomic pipeline to be compatible with tissues bioprinted directly onto microscope slides, eliminating embedding and sectioning steps.^33^ This capability is particularly powerful because it allows every aspect of the tissue to be controlled: cell types, number, and position, the inclusion of fluorescent reporters for ground-truth validation or real-time monitoring, and live-cell imaging to capture dynamic behaviors. All measurements can then be directly linked to transcriptome readouts, creating a highly tractable system for connecting spatial context to molecular state. Our data from this pipeline demonstrate that the arrangement of cells strongly influences their dynamic interactions, signaling environments, and transcriptional identities. Consistent with this, randomly deposited co-culture bioprints showed greater biological variability across replicates, reflecting the unpredictable spatial organization of local neighborhoods.

As with any emerging platform, several areas remain active targets for development. The platform supports reliable deposition across a broad range of cell types and biomaterials, with bioprinting success depending in part on the ability of deposited cells to rapidly adhere to the underlying substrate. Highly adherent cells readily bind at the point of deposition, whereas weakly adherent or suspension cells can be more challenging to position reproducibly. The precision of individual cell placement also creates opportunities for future automation, where closed-loop sensing could increase throughput without sacrificing spatial accuracy. Finally, the platform supports sequential cell deposition and hydrogel encapsulation to generate layered three-dimensional constructs. Extending this capability to increasingly complex architectures with single-cell z-resolution is an active direction of our ongoing work.

Taken together, these advances establish a generalizable framework for constructing and manipulating multicellular systems with single-cell spatial precision. By enabling spatial organization to be specified and directly linked to molecular and functional readouts, deterministic single-cell bioprinting provides a platform for causal investigation of how each cell contributes to the emergent behavior of the multicellular system. As spatial biology continues to reveal the importance of single-cell behavior in development, disease and therapeutic response, experimental systems with single-cell control may play an increasingly important role in translating descriptive spatial results into mechanistic understanding.

## Methods

### Human cell culture

Human myoepithelial cells were isolated from reduction mammoplasty tissue^34^ (kindly provided by Dr. William C. Hines lab, University of New Mexico) and cultured on a type I bovine collagen coating (PureCol, 100 µg/mL working concentration, Advanced Biomatrix, 5005) in M87 media. M87 media is a 1:1 mixture of Medium 171 (Gibco, M171500) and DMEM/F-12 (Gibco, 11330057) supplemented with 0.25% FBS (Cytiva, SH30396.03), 2mM L-glutamine (Gibco, 25030081), 0.1% w/v Albumax (Gibco, 11021037), 35 µg/mL bovine pituitary extract (Gibco, 13028014), 5 ng/mL epidermal growth factor (PeproTech, AF-100-15), 0.3 µg/mL hydrocortisone (Sigma, H0888), 0.5 ng/mL cholera toxin (Sigma, C8052), 7.5 µg/mL insulin (Sigma, I1882), 2.5 µg/mL apo-transferrin (Sigma, T1147), 0.1 nM oxytocin acetate salt (Sigma, O6379), 0.5 nM β-estradiol (Sigma, E2758), 5 nM Tri-iodothyronine (Sigma, T5516), and 5 µM isoproterenol (Sigma, I6504). Primary green fluorescent protein-expressing human umbilical vein endothelial cells (GFP-HUVEC, Angio-proteomie, cAP-0001GFP) were cultured in vascular endothelial growth factor endothelial growth medium (VascuLife VEGF, Lifeline Cell Technology, LL-0003). Primary bone marrow-derived mesenchymal stem cells (MSC, RoosterBio, MSC-003) were cultured in alphaMEM (Gibco, 12571071) supplemented with 10% MSC-qualified fetal bovine serum (FBS, Gibco, 12662029) and 1% penicillin-streptomycin (Gibco, 15140122). Primary mammary fibroblasts (ScienCel, 7630) were cultured in DMEM (Gibco, 11995073) supplemented with 10% FBS (Cytiva, SH30396.03) and 1% penicillin-streptomycin (Gibco, 15140122). Primary macrophages (Celprogen, 36070-01) were cultured in human macrophage cell culture medium with serum (Celprogen, M36070-01S). Human osteoblasts (ATCC, CRL-3602) were cultured in osteoblast growth medium (Cell Applications, 417-500). Cardiomyocytes (kindly provided by Dr. Christopher S. Chen’s lab, Boston University), were derived from human induced pluripotent stem cells (hiPSCs) via small molecule manipulation of the Wnt signaling pathway.^35,36^ iPSC-cardiomyocytes were cultured on Matrigel coated plates in RPMI supplemented with 1:50 B-27 (Gibco, 17504044) and 5% FBS (Gibco, 12662029). The culture was maintained until the cardiomyocytes resumed beating before harvesting for bioprinting. THP-1 derived macrophages were generated following a previously established protocol.^37^ Briefly, THP-1 were seeded into a 6 well plate at 2 x 10^5^ cells/well and treated with 20 ng/mL phorbol-12-myristate 13-acetate (PMA, Sigma, P8139) in RPMI 1640 (Gibco, 11875085) supplemented with 10% FBS (Gibco, 12662029). The media containing PMA was removed after 24 hours and cells were allowed to rest in fresh RPMI without PMA for 24 hours prior to use. Triple negative breast cancer MDA-MB-231 (ATCC, HTB-26), breast cancer MCF7 (Horizon Discovery), and neuroblastoma SH-SY5Y (ATCC, CRL-2266) were cultured in DMEM (Gibco, 11995073) supplemented with 10% FBS (Gibco, 12662029) and 1% penicillin-streptomycin (Gibco, 15140122). Prostate cancer PC3 (ATCC, CRL-1435) and monocyte THP-1 (ATCC, TIB-202) were cultured in RPMI 1640 (Gibco, 11875085) supplemented with 10% FBS (Gibco, 12662029) and 1% penicillin-streptomycin (Gibco, 15140122). Mammary epithelial MCF10A (Horizon Discovery, parental line; originating from ATCC CRL-10317) were cultured in DMEM/F12 (Gibco, 11330057) supplemented with 5% horse serum (Gibco, 26050088), 20 ng/mL epidermal growth factor (PeproTech, AF-100-15), 0.5 µg/mL hydrocortisone (Sigma, H0888), 100 ng/mL cholera toxin (Sigma, C8052), 10 µg/mL insulin (Sigma, I1882), and 1% penicillin-streptomycin (Gibco, 15140122). All cells were cultured at standard 37°C and 5% CO_2_. Unless otherwise stated, media was changed every 2-3 days and cells were passaged at 70% confluency using TrypLE Express (Gibco, 12605036).

#### Cell preparation for single-cell bioprinting

Cells were detached using TrypLE (Gibco, 12605036) express for the minimum possible time and pelleted at 300 g for 3-5 minutes. While bioprinting can be done label free, cells were either transduced to include a fluorescent reporter (iRFP or mVenus Histone 2B or tdTomato-9 EF1α (Takara Bio)) or fluorescently tagged using Hoechst (NucBlue Live ReadyProbes, Invitrogen, R37605) or general membrane cell tracker (Cellvue Claret Far Red, Sigma, MINCLARET; PKH26 Red Fluorescent Cell Linker, Sigma, PKH26GL; or PKH67 Green Fluorescent Cell Linker, Sigma, PKH67GL) following the manufacturer’s protocol to facilitate downstream analysis. Cells were then resuspended at 1 x 10^6^ cells/mL in a 1:1 mixture of complete cell culture medium and polyethylene glycol (PEG, 35,000 m.w., 30 mg/mL, Sigma, 81310).

### Biopixlar modifications

The microfluidic dispenser (Biopixlar, Fluicell, Mölndal, Sweden) configuration was adjusted so that the printhead was angled at 70° and retrofitted with a heating unit to maintain the chamber at 37 °C.^38^ A custom transparency overlay software was also developed to facilitate high precision patterning.^39^

#### Heating unit

Two 110 V, 100 W, 3.66 by 0.94 by 1.18 inch, ceramic air heater cartridges were used (ASIN: B01COPT7BG). A temperature control unit (model ITC-106VH) and solid state relay module (model SSR-40 DA) were purchased from Inkbird Tech C.L.. The temperature control unit and solid-state relay module were wired according to the documentation provided with the modules. Two ceramic heater cartridges were wired in parallel to the output of the solid-state relay module. The thermocouple was mounted onto the Biopixlar printhead holder near the tip and the heating modules were positioned on adjustable arms approximately 15 cm from the printing area. The controller was set to heat mode, with a threshold temperature of 37 °C, a control period of 4 seconds, a proportional band term of 30, an integration time of 30 seconds, and a derivative time of 8 seconds. Available at: https://github.com/ware-research/heater_feedback_controller/tree/v1.1

#### Bioprint map transparency overlay

A custom python program (Python v. 3.10.0) was developed to overlay a transparent image of the target bioprint map over the live microscope feed of the Biopixlar software.^39^ The software has been made available free of charge and can be accessed at: https://github.com/ware-research/transparency_overlay/tree/v.1.1

### General single-cell bioprinting setup

A 1.5 mg/mL collagen I (Gibco, A1048301) receiving substrate was prepared by adding 200 µL of collagen to the inner ring of a 35 mm, low walled petri dish (µ-Dish 35 mm low, Ibidi, 80156) and placed in the incubator for 30 minutes to crosslink. A 30 µm channeled PDMS printhead (FBPX-30, Fluicell) was primed by adding 25 µL of ultra-pure sterile water (Invitrogen, 10977015) to each well of the printhead and inserted into the priming pump (Fluicell) at 180 mbar for 3 minutes. Immediately prior to bioprinting, the water was removed from all waste collection wells and any wells which cells will be added to 25 µL of each cell suspension (1 x 10^6^ cells/mL in a 1:1 mixture of complete cell culture medium and 35,000 m.w. polyethylene glycol at 30 mg/mL) was added to the desired delivery wells. The printhead was loaded into the Biopixlar and prime 1 pre-set was executed (180 mbar delivery pressure to wells 1-4 simultaneously). 1.5 mL complete medium was added to the 35 mm dish containing the crosslinked collagen substrate and loaded into the printer. Following the completion of prime 1, the tip was gently wiped with the provided lint-free cloth (Fluicell) before lowering the printhead into the dish. Prime 2 pre-set was then executed to remove any bubbles in the fluidics (−220 mbar external and internal vacuums). The print surface was identified by dispensing a few cells onto the surface, bringing them into focus with the onboard microscope (LS620, Etaluma), and lowering the printhead next to the attached cells until a slight deflection was observed. If cells weren’t binding, the printhead was lowered at 5 µm increments. The cells used to identify the surface were removed by increasing the internal vacuum and aligning the dispensing channel with the cells. Bioprints were visualized in real-time during bioprinting using the onboard microscope (LS620, Etaluma). Fluorescent images of live cells, pre-labeled with a fluorescent tag prior to bioprinting, were acquired using either EVOS Fl Auto Imaging System (Life Technologies) or spinning disk confocal (Yokogawa CSU-X1 on Zeiss Axio Observer).

### Recirculation zone stability assessment

To assess the stability of the recirculation zone, fluorescein (200 µM in 1x PBS) was added into well 1 of the printhead and 1x PBS was added to wells 2-4. The printhead was lowered into a petri dish containing 1x PBS and a collagen substrate. Delivery pressure was set to either 60, 80, or 100 mbar, with non-delivery set to 0 mbar, and the internal and external vacuum settings were varied at 10 mbar increments from 0 to −100 mbar. The delivery pressure was applied for 10 seconds to observe the stability of the recirculation zone. If the area of the recirculation zone didn’t expand over the 10 second period, the recirculation was considered stable. A snapshot was taken at the 10 second mark before stopping the delivery pressure. This process was repeated across three different printheads (FBPX-30, Fluicell) to assess printhead-to-printhead variability.

### Maximum internal vacuum assessment

GFP-HUVECs were added to well 1 and 1x PBS was added to wells 2-4. The delivery pressure was set to either 60, 80, or 100 mbar with equal external vacuum and 0 mbar non-delivery pressure. The internal vacuum was varied at 10 mbar increments from 0 to −50 mbar to determine the range of internal vacuum settings that support cell dispensing. The delivery pressure of the cell containing well was applied for 10 seconds and cell dispensing (yes/no) was recorded. This process was repeated across three different printheads (FBPX-30, Fluicell), with *n* = 3 per printhead, to assess printhead-to-printhead variability.

### Minimum internal vacuum assessment

GFP-HUVECs were added to well 1 and GFP-HUVECs co-tagged with Hoechst (NucBlue Live ReadyProbes, Invitrogen, R37605) were added to wells 2-4. Delivery pressure was set to either 60, 80, or 100 mbar, with equal external vacuum and 0 mbar non-delivery pressure, and the internal vacuum was varied at 10 mbar increments from 0 to −40 mbar. GFP-HUVECs in well 1 were dispensed in a straight line for 10 seconds. If a Hoechst + GFP-HUVEC was dispensed, the internal vacuum setting was determined to be too low. This process was repeated across three different printheads (FBPX-30, Fluicell), with *n* = 3 per printhead, to assess printhead-to-printhead variability.

### Optimized single-cell bioprinting settings

Regardless of cell type, 1 x 10^6^ cells/mL in a 1:1 mixture of complete cell culture medium and polyethylene glycol (PEG, 35,000 m.w., 30 mg/mL, Sigma, 81310) was found to be suitable for single-cell deposition and binding onto a 1.5 mg/mL collagen I substrate. Optimized flow settings were 80 mbar delivery, 0 mbar non-delivery, −25 mbar internal vacuum, and −40 mbar external vacuum. For bioprints with cells positioned less than 25 µm away from one another, the external vacuum was dropped as low as −25 mbar to avoid disrupting the freshly deposited cells.

### Alternative receiving substrates

35 mm, low walled petri dishes (µ-Dish 35 mm low Grid-500, Ibidi, 80156) were prepared by coating the surface with one of the following materials: 0.1% w/v gelatin (from porcine skin, gel strength 300 Type A, Sigma, G2500) in sterile water and incubated at room temperature for 5 minutes, Matrigel (Corning, 354234) diluted 1:80 in DMEM/F12 and incubated at 37 °C for 1 hour, or 2 µg/mL Poly-L-Lysine (PLL, ScienCell, 0403) in sterile water and incubated at 37 °C for 1 hour. Excess solution was aspirated and replaced with 1.5 mL of the cells complete medium minus FBS. PLL coated dishes require two washes with sterile water before the addition of cell culture medium. To create three-dimensional stacks of collagen, the complete medium was aspirated from the dish following the completion of bioprinting. An additional 25 uL of collagen I was gently pipetted onto the surface and incubated at 37 °C and 5% CO_2_ for 15 minutes. Fresh medium was added to the dish and the bioprinting resumed on the new layer of collagen.

### Cell-cell interaction arrays

#### Arrays of varied cell arrangement

Triple negative breast cancer cells (MDA-MB-231) were stained with PKH26 Red Fluorescent Cell Linker (Sigma, PKH26GL) for general cell membrane labeling prior to bioprinting. Primary endothelial cells (GFP+ HUVEC) and triple negative breast cancer (TNBC) cells were bioprinted in a 4 x 4 array with each bioprint containing equal number of each cell type (8 endothelial and 8 TNBC) and 100 µm spacing between each cell. The spatial arrangement of the cells was either segregated (like cell types grouped half and half, 2×4), clustered (like cell types clusters in the corners, 2×2), or mixed (alternating cell types) to investigate the effect of spatial patterning on cell behavior. Bioprints were cultured in a 1:1 mixture of endothelial complete medium and TNBC complete medium. Once bioprinting was complete, samples were live-cell imaged on a spinning disk confocal (Yokogawa CSU-X1 on a Zeiss Axio Observer) with environment chamber at 37 °C and 5% CO_2_. Images were taken with fixed intervals of 10 minutes over 30 hours.

#### Arrays of varied cell composition

H2B-iRFP+ triple negative breast cancer (MDA-MB-231) were bioprinted in a 4 x 4 array of cells, with 100 µm spacing between each cell, either alone or in mixed (alternating configuration) with primary myoepithelial cells, primary mammary fibroblasts, or primary macrophages. Bioprints were cultured in M87 media and imaged at fixed intervals of 10 minutes for 24 hours on a spinning disk confocal (CrestOptics X-Light V3 on a Nikon Ti2) with environment chamber set at 35 °C and 5% CO_2_.

### Cell motion tracking, quantification, and analysis

Motion tracks were manually created in Imaris (Oxford Instruments, Version 10.2) by clicking the center of each cell per frame. Tracks were split when a cell division occurred and the daughter cell’s track began at the first frame it was visible. Raw *x* and *y* coordinates were exported for analysis.

#### Total distance traveled and normalized distance

Total distance traveled was calculated for each cell trajectory by summing the Euclidean distance between consecutive frames:

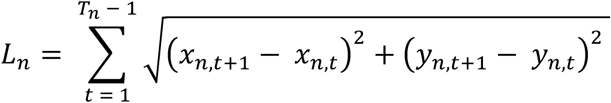

Where *T_n_* is the number of valid frames for cell *n*, and (*x_n_*_,*t*_, *y_n_*_,*t*_) are the raw *x*, *y* coordinates at frame *t*.

To account for cells that traveled out of the field of view or divided, which would truncate track length sums, we normalized distanced traveled by the number of consecutive frames observed:

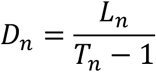

Where *D_n_* is the distance traveled per visible frame for cell *n*, and (*T_n_* − 1) is the number of consecutive frame intervals over which the cell was observed.

#### Mean squared displacement

The time averaged mean squared displacement (TASD)^40^ was calculated for each cell trajectory using the following equation:

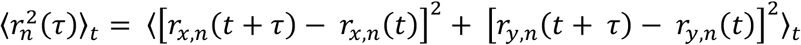

Where *r_x_*_,*n*_(*t*) and *r_y_*_,*n*_(*t*) are the *x* and *y* coordinates of cell *n* at time *t*, and 〈⋅〉*_t_* denotes the average taken over all time points for which a corresponding position at *t* + *τ* exists. Cells were imaged at fixed interval Δ*t*, so lag times occurred at discrete steps *τ* = *k*Δ*t*, where *k* is the lag in frames. TASD values 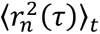 were computed for each cell and then averaged across all cells belonging to a given experimental group. The standard error of the mean (SEM) at each *τ* was calculated across cells contributing measurements at that lag.

To assess how cell migration scaled over time, we examined the power-law relationship^40^

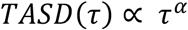

Where *α* is the anomalous diffusion exponent describing the mode of migration. Subdiffusive motion (0 ≤ *α* < 1) reflects restricted or confined movement, *α* = 1 corresponds to Brownian (random) motion, and *α* > 1 indicates superdiffusive or persistent migration. For each cell type and spatial arrangement, *α* was extracted by fitting the linearized form:

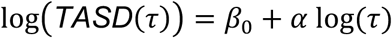

Only finite TASD values greater than zero were retained, TASD values within a defined lag-time window of 50-500 minutes were used, and a minimum of three points was required for a valid fit. The slope of the regression yielded *α*, and the coefficient of determination (*R*^2^) was used to assess the quality of the fit.

#### Net displacement

Net displacement for each cell was calculated using the following equation:

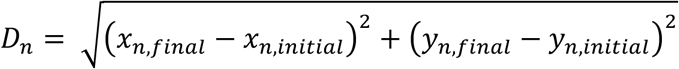

Where (*x_n_*_,*initial*_, *y_n_*_,*initial*_) is cell the starting position of cell *n* and (*x_n_*_,*final*_, *y_n_*_,*final*_) is the final position of that cell.

#### Center of mass displacement

The center of mass (COM) of the HUVEC population was calculated separately for each bioprinted array by averaging the *x* and *y* coordinates of all tracked cells. For each replicate at time *t*, COM was defined as:

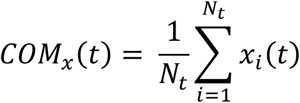

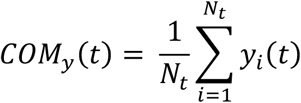

Where *N_t_* is the number of cells contributing positional measurements at time *t*. To quantify population-level shifts over time, COM trajectories were centered such that the initial position served as the origin:

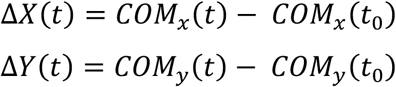

For each spatial arrangement, Δ*X*(*t*) and Δ*Y*(*t*) curves were averaged across independent bioprinted arrays (three arrays per arrangement), and the standard error of the mean (SEM) was calculated at each time point. For each array, a time averaged COM displacement was then calculated along *X* and *Y* by averaging Δ*X*(*t*) and Δ*Y*(*t*) over the full 30 hour imaging period.

#### Distance to nearest neighbor

For each MDA-MB-23, the distance to its nearest non-MDA-MB-231 neighbor was computed:

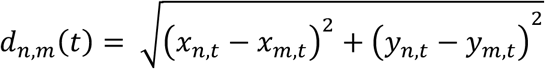

Where (*x_n_*_,*t*_, *y_n_*_,*t*_) are the coordinates of MDA-MB-231 cell *n* and (*x_m_*_,*t*_, *y_m_*_,*t*_) are the coordinates of non-MDA-MB-231 cell *m* in the same time frame *t*.The nearest neighbor distance for cell n was then defined as:

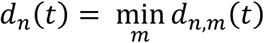

For each experimental group (TNBC + Myoepithelial, TNBC + Fibroblast, or TNBC + Macrophage) and time point, nearest neighbor distances were averaged across all MDA-MB-231 (TNBC) cells:

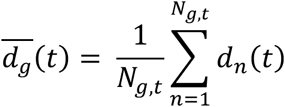

Where *g* is the experimental group, 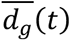 is the mean nearest neighbor distance at time *t*, and *N*_g,*t*_ is the number of MDA-MB-231 cells contributing measurements.

#### Migration territory

To quantify the two-dimensional territory explored by migrating cells, trajectories were first translated so that each track began at the origin. For each cell *n*, the recorded positions (*x*_(*n*,*t*)_, *y*_(*n*,*t*)_) were converted to relative coordinates

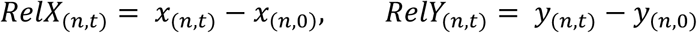

such that (*RelX*_(*n*,0)_, *RelY*_(*n*,0)_) = (0,0). For each sample, the spatial territory explored was defined by the area of the convex hull surrounding all relative coordinates of the cell type of interest. Convex hulls were generated for each sample using the Qhull algorithm (convhulln, geometry package).

#### Travel consistency

Travel consistency, also known as the consistency index, was calculated for each cell *n* as the ratio of net displacement to total distance traveled:

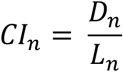

Where *D_n_* represents the Euclidean distance between the first and last position of the cell trajectory, and *L_n_* denotes the cumulative Euclidean distance traveled between consecutive frames along the trajectory.

### Breast cancer biopsy recreation

To replicate a tumor biopsy region of interest, a breast cancer biopsy previously annotated using imaging mass cytometry was selected.^25^ A region containing a breast duct with surrounding stroma was selected and imported into ImageJ. The image was converted to an 8-bit image and threshold to segment each cell. Cells that were incorrectly fused were manually separated by drawing an eraser line. The particles were then analyzed and any cell with an area less than 254.47 µm^2^ (18 µm diameter, ∼the average size of our detached cells) was removed. The resulting masked image was manually re-colored using Procreate (Savage Interactive) to return the annotations. A 3D bioprint map model was then created using Fusion 360 (Autodesk CAD) by placing 18 µm diameter spheres in the center of each cell of the mask. In regions where cells had been removed due to size limitation, spheres were evenly distributed to fill in the void. The resulting 3D model was exported as the bioprint map with the X,Y location of each cell. Using our custom overlay software^39^, the bioprint map was matched using mammary epithelial cell MCF10A, triple negative breast cancer cell MDA-MB-231, primary mammary fibroblasts, primary mesenchymal stem/stromal cells, and THP-1 derived macrophages. The bioprint was cultured in a 1:6 mixture of mammary epithelial MCF10A media and DMEM supplemented with 10% FBS and 1% penicillin-streptomycin. Live-cell imaging was conducted using a widefield fluorescent microscope (Nikon Ti2), acquiring an image every 10 minutes over 24 hours, with environment chamber at 37 °C and 5% CO_2_.

### Biopsy replica fidelity analysis

The fluorescence image of the resulting bioprints were imported into Fusion360 (Autodesk CAD). 18 µm diameter spheres were placed in the center of each cell of the image, following the same approach used to generate the print map. Fidelity analysis was performed using one full-field image of the biopsy replica and three smaller subregions sampled from three additional prints. The resulting 3D models of the bioprints and the original bioprint map 3D model were exported as .OBJ files and imported into CloudCompare (v2.13.2). A cloud to mesh (C2M) distance comparison was computed using the default settings (octree level: auto; absolute distances). The points were visualized with a histogram, and a Gaussian distribution was fitted to determine bioprint fidelity. Values reported as average ± standard deviation.

### Breast tumor microenvironment model

A pre-invasive breast cancer model was fabricated by first bioprinting 200 µm diameter rings of mammary epithelial cells (MCF10A) onto a collagen substrate in M87 medium. The bioprints were returned to the incubator while the remaining cell types were harvested. Triple negative breast cancer cells (MDA-MB-231) were deposited within the epithelial ring and primary mammary fibroblasts were patterned around the epithelial rings and throughout the stromal compartment. A thick fibroblast box “wall” was bioprinted around the entire bioprint to serve as physical boundaries to promote cells staying within the region instead of migrating to fill in the vast open spaces of the 35 mm petri dish. To simulate standard co-culture approaches, mammary epithelial (MCF10A), triple negative breast cancer (MDA-MB-231), and mammary fibroblasts were mixed at a 2:1:5 ratio respectively and loaded into a single well of the printhead. Cells were deposited in a random distribution, and no box perimeter was bioprinted. Immediately following bioprinting, live-cell imaging began on a widefield fluorescent microscope (Nikon Ti2) with environment maintained at 37 °C and 5% CO_2_. Images were acquired every 10 minutes for 24 hours.

### Immunostaining

Bioprints were washed three times with 1x PBS and fixed using 10% neutral buffered formalin (Epredia, 22-110-873) for 30 minutes at room temperature. Cells were permeabilized using 0.1% Triton X-100 (Fischer Scientific, BP151) solution in PBS for 15 minutes at room temperature. The samples were then blocked using 1.5% bovine serum albumin (BSA, Fischer Scientific, BP1600) in PBS for 1 hour at room temperature. Primary antibodies pan cytokeratin (PanCK, 1:100, OriGene, CF190321) and vimentin (VIM, 1:200, Novus Biologicals, NBP131327) were added in 0.15% BSA and incubated at 4 °C overnight. Primary antibodies were removed, and the samples were washed three times with PBS before adding secondary antibodies (1:250, goat anti-mouse Alexa Fluor 555, Invitrogen, A-21422; goat anti-rabbit Alexa Fluor 647, Invitrogen, A-21244). Secondary antibodies were incubated at room temperature for 2 hours, removed, and samples washed twice with 0.1% Tween 20 (Thermo Scientific, J20605.AP). Nuclei were labeled with DAPI (NucBlue Fixed Cell ReadyProbes, Invitrogen, R37606) in PBS for 30 minutes at room temperature. Samples were then imaged using a spinning disk confocal (Yokogawa CSU-X1 on Zeiss Axio Observer).

### Spatial transcriptomics

The Visium HD Spatial Gene Expression assay (10x Genomics, Human Transcriptome, 6.5 mm) was modified as we previously reported^33^ to enable analysis of planar cell cultures without sectioning and embedding. Briefly, a standard microscope glass slide was sterilized in 70% ethanol, rinsed with ultrapure sterile water, and coated with type I bovine collagen solution (PureCol, 100 µg/mL working concentration, Advanced Biomatrix). Cells were bioprinted directly onto the treated slide submerged in M87 medium to create the following groups: spatially defined tumor microenvironment, randomly deposited co-culture, and monoculture controls. The spatially defined tumor microenvironment samples were created by bioprinting four tdTomato9-EF1α+ mammary epithelial (MCF10A) rings (200 µm in diameter), with H2B-iRFP+ breast cancer (MCF7) bioprinted in the center and H2B-mVenus+ primary mammary fibroblasts surrounding the rings. Co-culture samples were fabricated by mixing the same ratio of cells as the patterned sample (1:2:5 MCF7:MCF10A:FB), loading into a single well of the printhead, and depositing over an equivalent area and density as the patterned samples. Single cell type controls were bioprinted onto one of the capture areas to serve as internal references. Samples were cultured for 24 hours. Immediately prior to fixation, a fluorescent image was taken to serve as a ground truth cell type identifier, as each cell type was transduced with a unique fluorescent reporter. Samples were then fixed with 4% paraformaldehyde (PFA) for 30 minutes at room temperature and permeabilized in 100% methanol on ice for 1 hour. H&E staining, imaging, destaining, and decrosslinking were performed according to the Visium HD FFPE Tissue Preparation Handbook (CG000684, Rev A). The manufacturer’s protocol (Visium HD Spatial Gene Expression User Guide CG000685, Rev B) was followed thereafter to generate the spatial gene expression libraries. The library was sequenced by NovoGene on an Illumina NovaSeq X Plus with a depth of approximately 400 million reads per sample. FASTQ files, H&E images, and spatial metadata were processed using Space Ranger v3.1.3 with the GRCh38-2020-A human reference genome and Visium human transcriptome probe set v2.0.

### Spatial transcriptomics data processing

Space Ranger outputs for each Visium HD capture area were imported into R (v4.4.2) using the Load10X_Spatial function in Seurat (v5) at an 8 µm bin size, and each object was assigned a unique capture area identifier. Custom metadata were curated using Loupe Browser (10x Genomics, v8) as previously described.^33^ Because the versions of Loupe Browser and Space Ranger available at the time of experimentation did not allow for the simultaneous alignment and processing of an IF image and an H&E image, our custom transparency overlay software^39^ was used to superimpose the fluorescent image of each sample taken right before fixation onto the Loupe Browser image. The custom group and paintbrush tools in Loupe were then used to manually annotate each barcode based on the fluorescence and cell morphology (H2B-iRFP+ MCF7, tdTomato9-EF1α+ MCF10A, and H2B-mVenus+ mammary fibroblasts). The annotated barcodes were exported as a .csv file and integrated into the Seurat object metadata. The custom group and rectangle selection tool in Loupe Browser was also used to assign sample ID and condition (Print Pattern) to each barcode. Datasets from the two capture areas were then merged, and Raw counts were log-normalized (LogNormalize, scale factor = 10,000) to generate the normalized data layer used for downstream analyses.

#### Gene ontology

For pathway-level comparisons between spatially defined and co-culture bioprints, barcodes belonging to each bioprint replicate were aggregated, and differential expression was performed between spatially defined and co-culture conditions in Seurat (Wilcoxon test, Benjamini–Hochberg correction). Genes significantly upregulated in each condition (adjusted P < 0.05) were converted to Entrez IDs and analyzed using clusterProfiler^41^ for Gene Ontology Biological Process over-representation (enrichGO, OrgDb = org.Hs.eg.db, ontology = “BP”). The top enriched terms for spatially defined and co-culture were visualized as mirrored −log10(adjusted P) bar plots.

### Single-spot gene set enrichment analysis (ssGSEA)

To quantify pathway activity at single-spot resolution, we performed ssGSEA using the Escape^42^ (runEscape, method = “ssGSEA”) on log-normalized expression values. Cancer hallmark^26^ gene sets (MSigDB collection H) were retrieved via msigdbr^43,44^, and a curated panel of 19 cancer-relevant pathways (E2F targets, G2M checkpoint, MYC targets v1 and v2, P53, TGFβ signaling, apoptosis, unfolded protein response, DNA repair, epithelial mesenchymal transition, apical junction, oxidative phosphorylation, glycolysis, interferon gamma response, complement, inflammatory response, TNFα via NF-κB, IL6–JAK–STAT3 signaling, and angiogenesis) were used for scoring. ssGSEA scores were computed independently for each spatial barcode, z-scaled across the dataset, and aggregated within each cell line (MCF10A, MCF7, fibroblast) and culture condition (spatially defined, co-culture, and monoculture control). Differences in pathway activity were evaluated using Wilcoxon rank-sum tests, including comparisons to mono-culture controls.

### Ligand-receptor analysis

Ligand–receptor interactions were inferred using LIANA^27,45^ (version 1.6.1; Python) with the rank_aggregate framework and consensus ligand-receptor resource. Cell type barcode annotations were used to define sender and receiver populations, and interactions were filtered using an expression proportion threshold of 0.1 with 1,000 permutations (remaining parameters default). Ranked interaction outputs were exported for downstream visualization in R. For display, rank-based LIANA metrics were transformed such that higher values indicate stronger or more specific interactions (e.g., 1 − magnitude_rank and 1 − specificity_rank).

To evaluate how cell–cell signaling differed between spatial arrangements, ligand–receptor results were aggregated by sender–receiver pair to construct directed signaling networks for spatially defined and co-culture bioprints. Networks were visualized as chord diagrams (Circlize^46^; R), with edge weights corresponding to the number of ligand–receptor pairs connecting each source–target pair. Sender–receiver rewiring was quantified by computing the change in ligand–receptor pair counts for each directed cell-type pair (Δ pairs = Spatially defined – Co-culture) and visualized as a dot plot, with color encoding the directionality of rewiring and point size indicating its magnitude.

To assess changes at the level of individual signaling interactions, we examined the highest-ranked ligand–receptor pairs within selected sender–receiver interactions by comparing condition-specific LIANA interaction scores. For each directed cell-type pair, non-overlapping top-ranked ligand–receptor pairs in spatially defined and co-culture configurations were identified and visualized based on the difference in interaction strength (Spatially defined – co-culture), highlighting ligand–receptor interactions preferentially utilized under each spatial arrangement.

To link inferred cell–cell communication with downstream regulatory programs in receiver populations, ligand–receptor interactions identified by LIANA were integrated with transcription factor and pathway activity inference using decoupler^28^ (version 2.1.2; Python). For each sender–receiver cell type pair, TF activities in the receiver cells were estimated from gene expression using the CollecTRI^29^ regulatory network and the ULM method, while pathway activities were inferred using the PROGENy^30^ model and the MLM method (human, top 100 target genes per pathway). Receptor–TF and TF–pathway associations were defined by Spearman correlation across receiver cells, and network edge weights were normalized across spatial conditions to enable direct comparison of inferred signaling architectures.

### Statistical analysis

Differential gene-expression and Gene Ontology Biological Process enrichment analyses were evaluated using adjusted P values with Benjamini-Hochberg correction. Pairwise two-sided Wilcoxon rank-sum tests were used to compare ssGSEA z-scores between spatially defined and standard co-culture conditions within annotated cell populations, with Benjamini-Hochberg correction for multiple comparisons. Ligand-receptor interactions were inferred using LIANA and summarized using method-specific ranks, scores and ligand-receptor pair counts. Associations linking receptor, transcription factor and pathway activities were evaluated using Spearman correlation across receiver cells. Cell migration and spatial organization metrics, including distance traveled, normalized distance, net displacement, center-of-mass displacement, convex hull area and confinement index, were compared across experimental conditions using one-way ANOVA with Tukey correction for multiple comparisons when three groups were analyzed. For comparisons involving four microenvironment composition conditions, including TNBC, TNBC+Myoep, TNBC+FB and TNBC+Mac, global differences were assessed using Welch’s one-way ANOVA to account for unequal variances, followed by pairwise Welch t-tests with Benjamini-Hochberg correction. The anomalous diffusion exponent α was estimated by linear regression of log-transformed time-averaged mean squared displacement over finite values greater than zero within the defined 50-500 min lag-time window, with at least three points required for a valid fit.

Data are shown as individual observations with violin/boxplot overlays at the per-cell or per-track level, and as mean ± SEM for time-resolved summary trajectories, including center-of-mass displacement, mean squared displacement and nearest-neighbor distance. Statistical significance was defined as adjusted P < 0.05 unless otherwise stated. Complete statistical reporting for each analysis, including sample sizes, summary statistics, test details, confidence intervals where applicable, adjusted P values and effect-size estimates, is provided in the Source Data.

## Materials availability

Printer modifications have been made available free of charge on GitHub: https://github.com/ware-research/heater_feedback_controller/tree/v1.1 and https://github.com/ware-research/transparency_overlay/tree/v.1.1.

## Data availability

The spatial transcriptomic dataset generated during this study has been deposited in the National Center for Biotechnology Information (NCBI) Gene Expression Omnibus and is accessible through GEO Series accession number GSE315233: https://www.ncbi.nlm.nih.gov/geo/query/acc.cgi?acc=GSE315233. Additional source data are provided on GitHub https://github.com/HaylieHelms/2DscBioprinting2 and Zenodo https://doi.org/10.5281/zenodo.18563242.

## Code availability

The code used to analyze the cell-cell interaction arrays and spatial transcriptomics data is available at: https://github.com/HaylieHelms/2DscBioprinting2 and has also been deposited at Zenodo: https://zenodo.org/records/18563242.

## Acknowledgements

The authors thank Michael McLellan of Christopher Chen’s lab for generating the hiPSC derived cardiomyocytes, Ryan Gordon for sharing the prostate cancer cell line PC3, Jens Kreth for sharing the monocyte cell line THP-1, Avathamsa Athirasala for generating the transduced MCF10A cell line, Rawan Makkawi for their collaboration to generate the histone 2B lentiviral reporters used to transduce the MCF7, MDA-MB-231 and fibroblasts, and Jason Ware for building the printer modifications. This work was supported by funding from the Cancer Early Detection Advanced Research Center, Knight Cancer Institute, Oregon Health and Science University (Full 2023-1745) awarded to H.R.H. and L.E.B, OHSU Silver Family Innovation Award awarded to L.E.B, and the M.J. Murdock Charitable Trust. Institutional support was provided by the OHSU Knight Cancer Institute NCI Cancer Center Support Grant (P30CA069533) for the Advanced Light Microscopy Core (RRID:SCR_009961). L.E.B. also acknowledges support from the National Institute of Dental and Craniofacial Research (R01DE035326), Knight Cancer Institute Pilot Award, and Cancer Early Detection Advanced Research Center (Full 2023-1719). H.R.H. acknowledges support from the International Alliance for Cancer Early Detection (ACED) PhD Fellowship and Achievement Rewards for College Scientists (ARCS) Foundation Oregon. A.E.D. acknowledges support from National Institutes of Health, Office of Research Infrastructure Programs (K01OD031811). E.M.L. acknowledges support from a Research Scholar Grant (RSG-22-060-01-MM) from the American Cancer Society (https:doi.org/10.53354/pc.gr.153686) and an ACED Project Award (ACNCR1743). The authors also thank the Cell, Developmental and Cancer Biology Summer Research Internship program at the Knight Cancer Institute, Oregon Health and Science University for supporting K.A.O. The International Alliance for Cancer Early Detection (ACED) is a partnership between Cancer Research UK, Dana Farber Cancer Institute, The University of Manchester, the German Cancer Research Center (DKFZ), University College London, Knight Cancer Institute at OHSU, and The University of Cambridge.

## Author contributions

H.R.H. designed the research with L.E.B; acquired funding; conceptualized printer modifications; performed the experiments; developed the spatial transcriptomics assay modifications; analyzed the data; and wrote the manuscript. K.A.O. worked with H.R.H. to create the patient matched bioprint map, assisted during bioprinting, and contributed to cell tracking. A.E.D. and E.M.L advised experimental design and imaging; and reviewed and edited the manuscript. L.E.B. conceived the research; acquired funding; supervised the work; and edited the manuscript.

## Competing interests

L.E.B is a co-founder of and has an equity interest in RegendoDent Inc. L.E.B is named on patents that have been licensed by RegendoDent Inc. His interests were reviewed and managed in accordance with the OHSU institutional conflict of interest policies. E.M.L. is an inventor on bioprinting technology that has been optioned to Organovo, Inc. by OHSU. This potential conflict of interest has been reviewed and managed by OHSU.

## Notes

### Summary of Updates

Updated manuscript text and figures 1-4.

