## Supplementary Figures for "A Single-Cell Bioprinting Method to Reconstruct Native Cellular Microenvironments with Subcellular Resolution"

### SUPPLEMENT FIGURES

| Instrument | Reproducible single-cell deposition? | | Smallest single-cell spacing demonstrated | | Max cell types/ populations used in a print | Demonstrate Native Tissue Reconstruction? | Citation |
|---|---|---|---|---|---|---|---|
| Laser Bioprinting | | | | | | | |
| NGB-R, Poietis | No | Missing spots and doublets+ 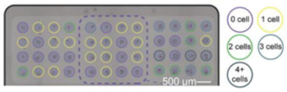 | 500 µm | Distance measured between spots, variable cell placement within each spot 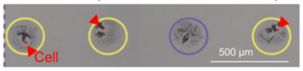 | 2<br>HUVEC and hMSC | No | Bosmans et al. [1] |
| Custom | Yes | One cell per target 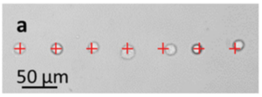 | < 20 µm | Inconsistent spacing and control 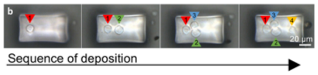 | 2<br>hMSC and hTSPCs | No | Zhang, J. et al. [2] |
| Inkjet Bioprinting | | | | | | | |
| Custom | Yes | One cell per spot, no missing spots 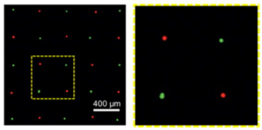 | 400 µm | 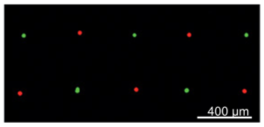 | 3<br>NIH 3T3 in three colors | No | Zhang, P. et al. [3] |
| JetLab II, MicroFab Technologies | No | Missing spots and doublets+ 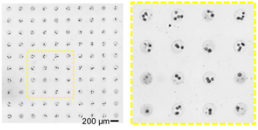 | 200 µm | Distance measured between spots, variable cell placement within each spot 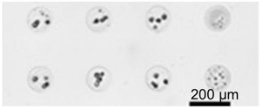 | 3<br>NIH3T3 in three colors | No | Park et al. [4] |
| Microfluidic Bioprinting | | | | | | | |
| Biopixlar, Fluicell | Yes | One cell per spot, no missing spots 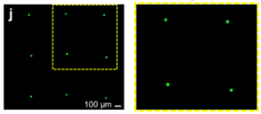 | 500 µm | 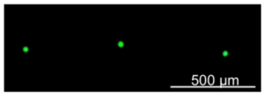 | 2<br>HaCaT and A431 | No | Jeffries et al. [5] |

**Table S1. Benchmarking of existing bioprinting platforms for deterministic single-cell deposition and native density spatial patterning.**
Images adapted with permission.

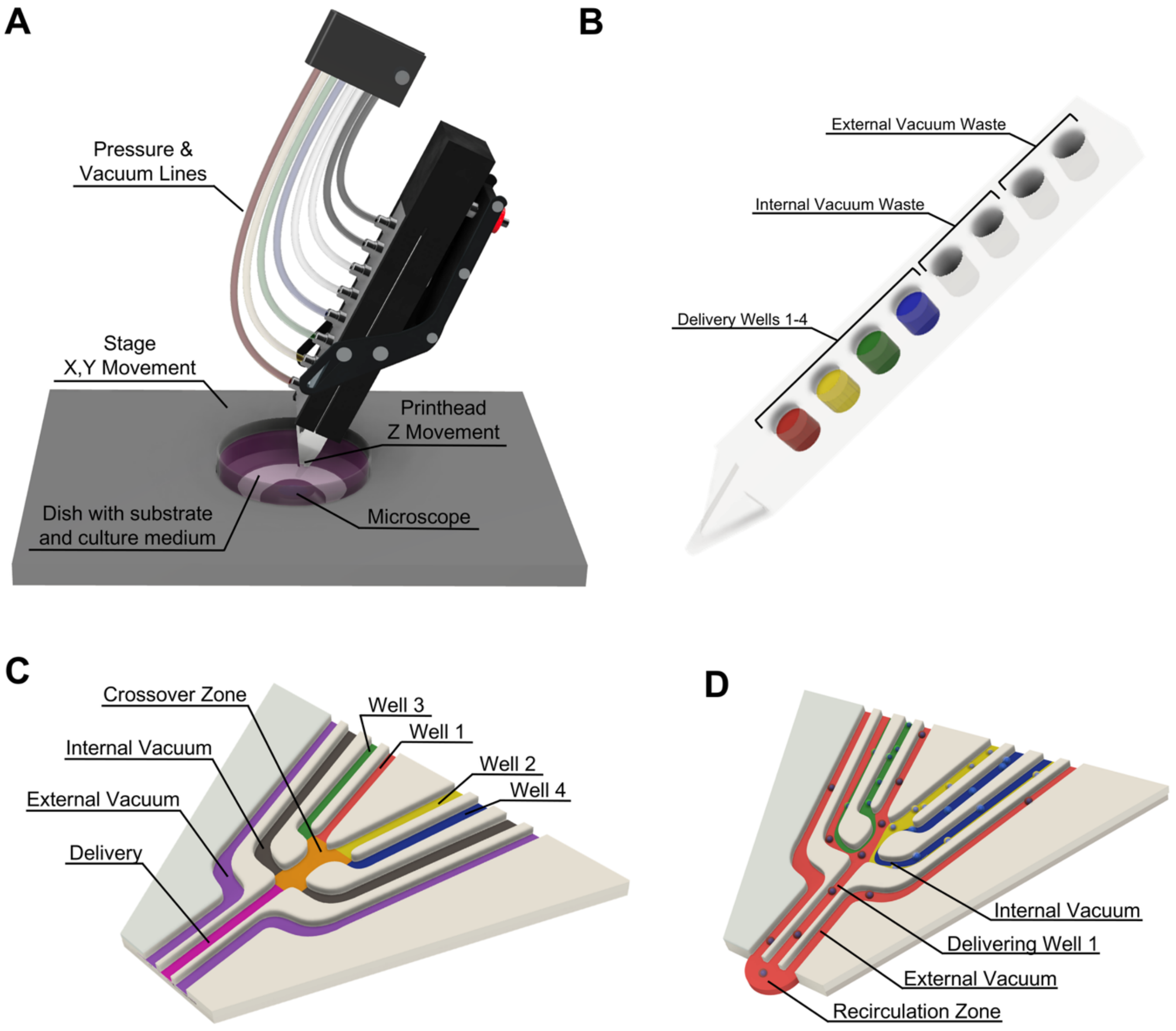


**Figure S1. Microfluidic dispensing system**
(A) Components of the microfluidic system.
(B) Printhead made from PDMS containing eight independent wells, four for delivery and four for waste collection.
(C) Tip of the printhead with annotated fluid flow regions.
(D) Tip of the printhead illustrating the dispensing of well 1.

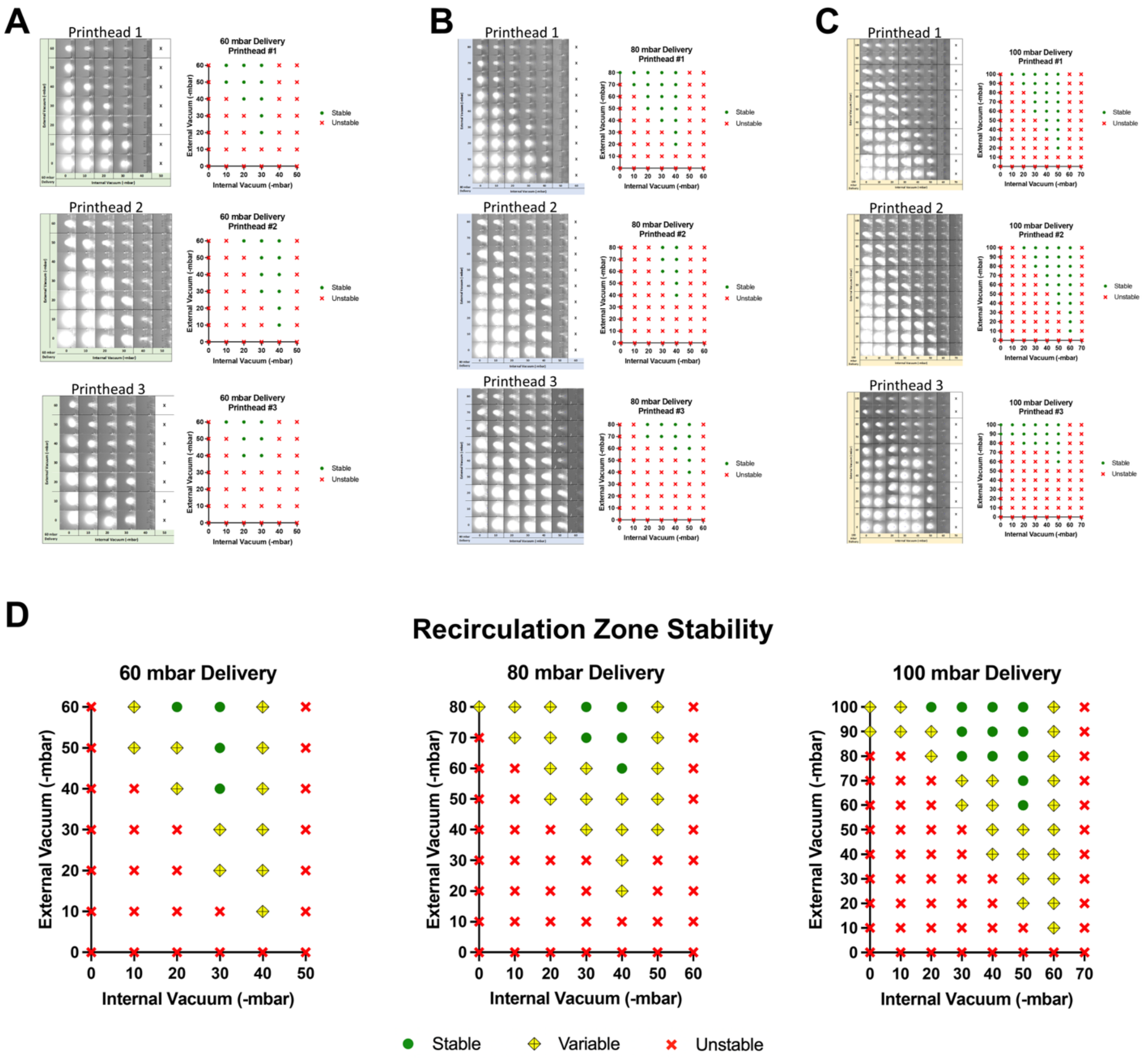


**Figure S2. Fluidics optimization for a stable recirculation zone at 60, 80, and 100 mbar delivery pressure**
(A-C) Recirculation zone visualization and stability across 3 printheads at A) 60 mbar B) 80 mbar and C) 100 mbar delivery pressure.
(D) Recirculation zone stability summary, n = 3.

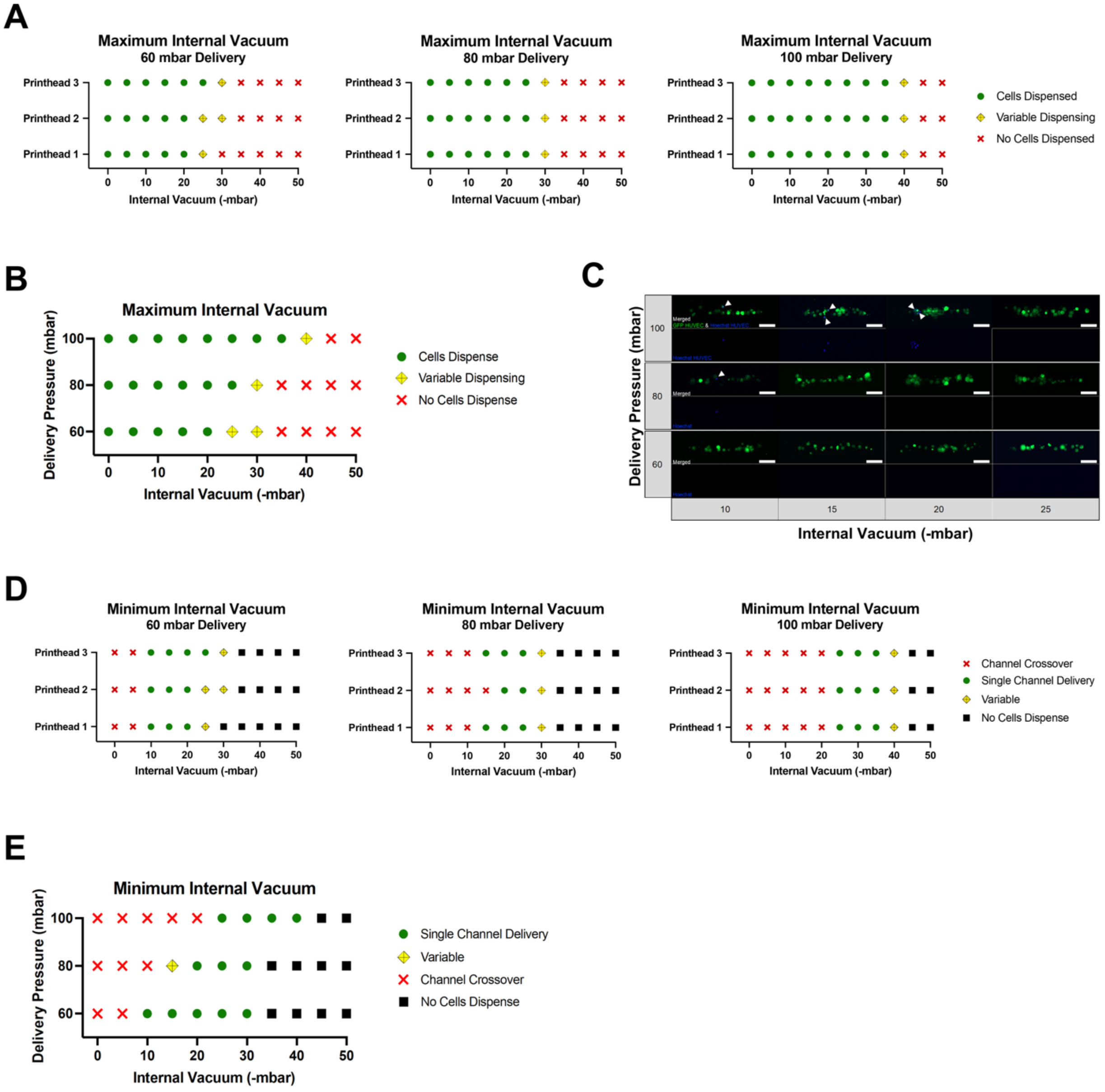


**Figure S3. Vacuum optimization for single-cell bioprinting at 60, 80, and 100 mbar delivery pressure**

(A-B) Maximum internal vacuum which allows to be dispensed. Variable indicates failure in at least one of the tests. (A) *n* = 3. (B) Study summary. *n* = 9.

(C-E) Minimum internal vacuum investigation to prevent unwanted cell dispensing. (C) Delivery of a Hoechst positive HUVEC indicates crossover from another delivery well and the internal vacuum setting is too low. White arrows indicated Hoechst+ HUVEC. Scale bars, 100 µm. (D) *n* = 3. E) Study summary. *n* = 9.

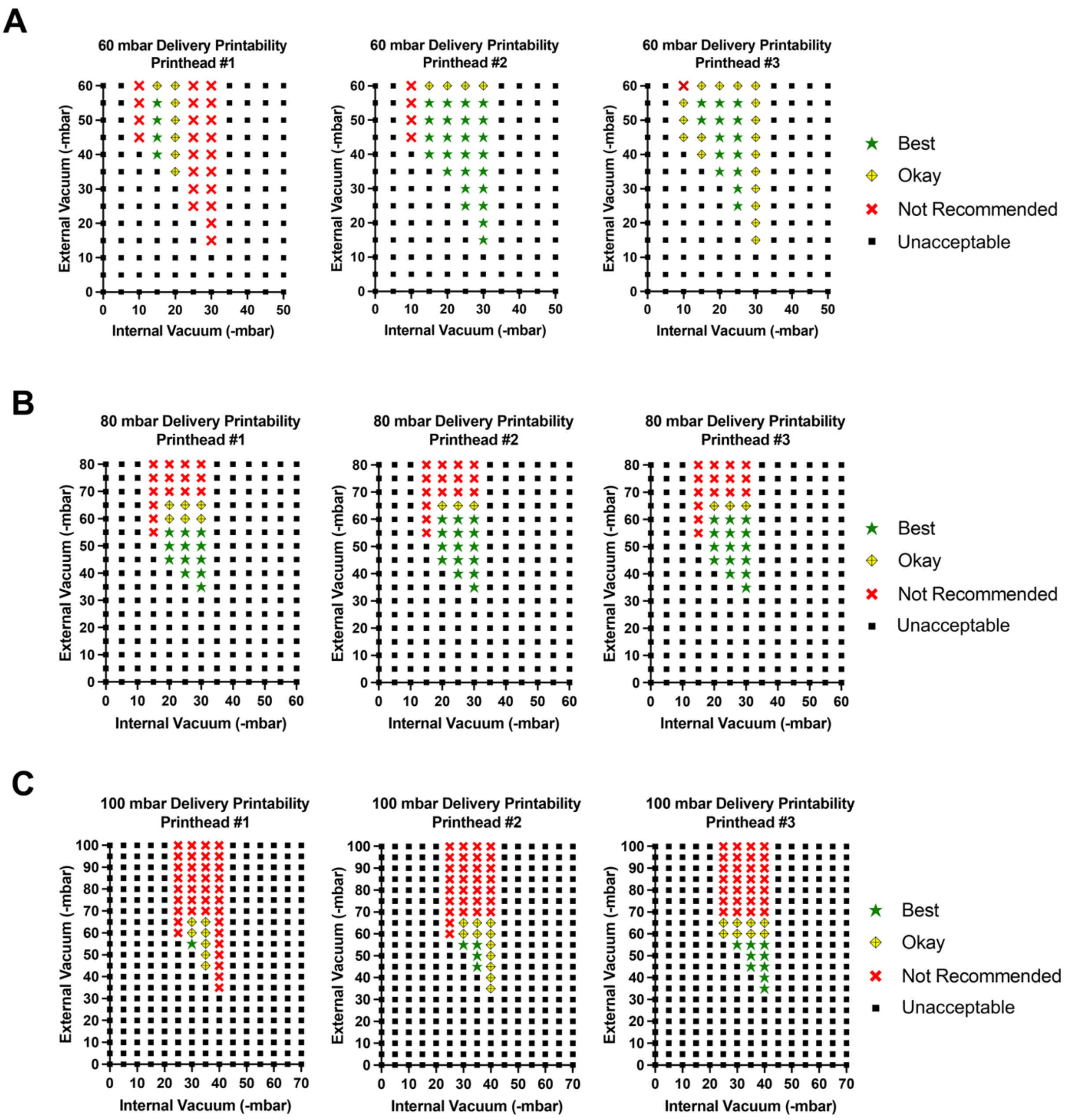


**Figure S4. Overall printability assessment**
Best: Conditions that meet optimization criteria and yield high-fidelity single-cell printing. Okay: Conditions that meet optimization criteria and support single-cell printing with moderate performance. Not Recommended: Conditions that satisfy optimization criteria but result in poor single-cell printing performance, such as inconsistent delivery. Unacceptable: Conditions that fail to meet one or more of the optimization criteria, including recirculation zone stability, cell delivery (max internal vacuum), and controlled multiplexing (min internal vacuum).
(A) 60 mbar delivery pressure. *n = 3.*
(B) 80 mbar delivery pressure. *n = 3.*
(C) 100 mbar delivery pressure. *n = 3.*

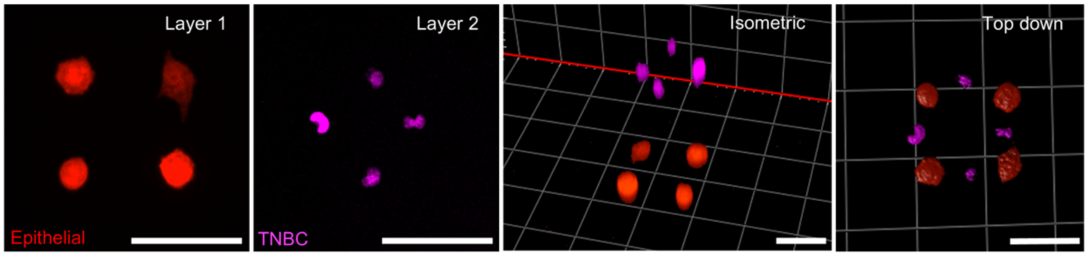


**Figure S5. 3D single-cell bioprint via collagen layering**
Scale bars, 50 μm.

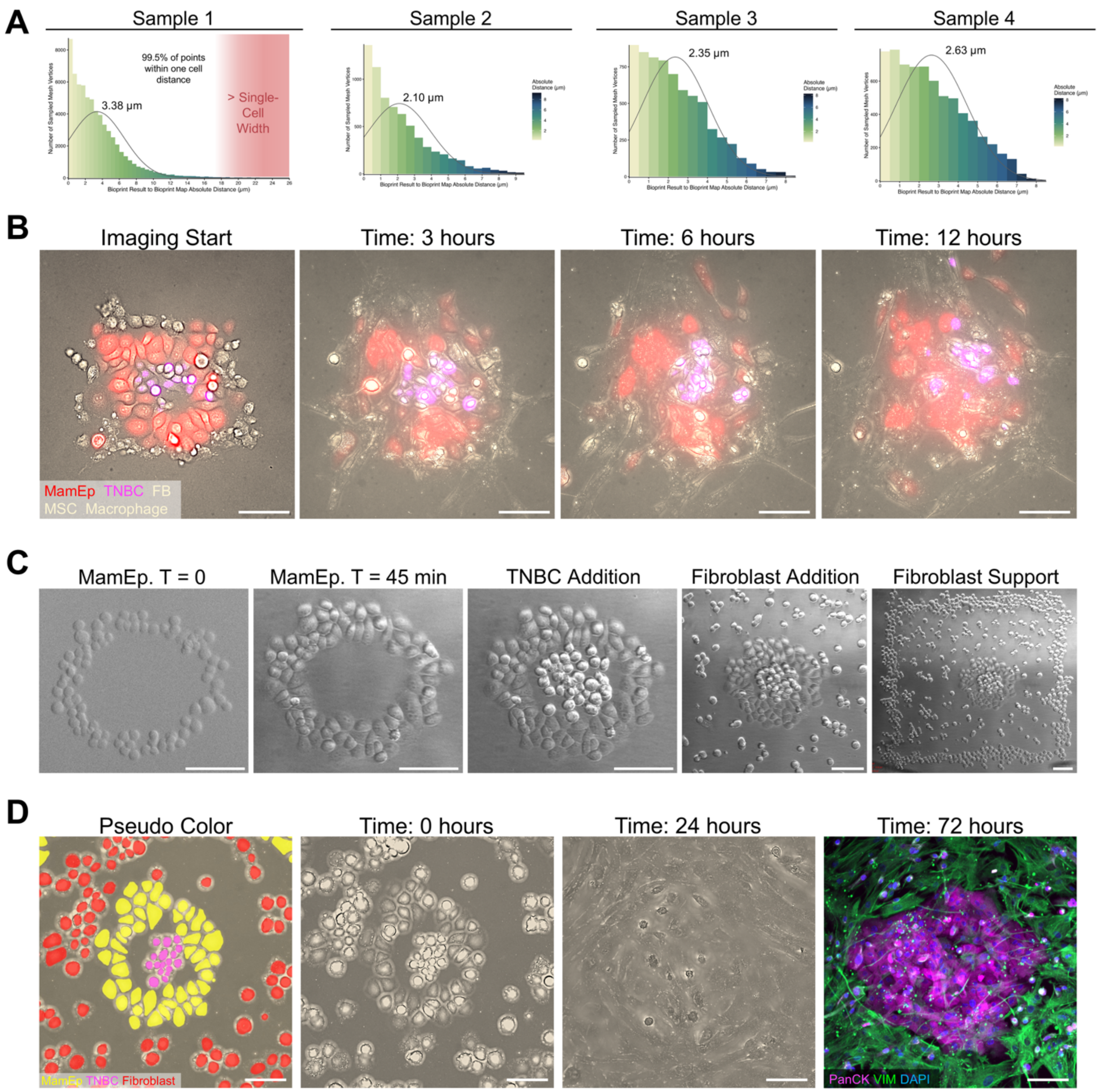


**Figure S6. Single-cell bioprinted breast tumor microenvironments**

(A) Bioprint fidelity measurements. Distribution of absolute distances between the bioprint result and reference bioprint map, computed from mesh vertices to the reference mesh surface. Single-cell diameter cutoff set at 18 µm.

(B) Live-cell imaging of high-fidelity replica at imaging time 0, 3, 6, and 12 hours. Scale bars, 100 µm.

(C) Pre-invasive breast cancer model fabrication. First MCF10A cells are bioprinted in a ring. The bioprint is returned to the incubator while the other cell types are harvested and prepared for bioprinting (30-45 minutes). MDA-MB-231 are added to the center of the ring and fibroblasts are patterned around it. To facilitate dense tissue formation, a thick box of fibroblasts is patterned around the tumor microenvironment to contain the bioprint and support growth as a feeder layer. Scale bars, 100 µm.

(D) Model pseudo colored to identify cell types: mammary epithelial (MCF10A), triple negative breast cancer (MDA-MB-231), and mammary fibroblasts. Live-cell images at 0 and 24 hours, followed by immunofluorescence staining at 72 hours for pan cytokeratin (PanCK), vimentin (VIM), and nucleus (DAPI). Scale bars, 100 µm.

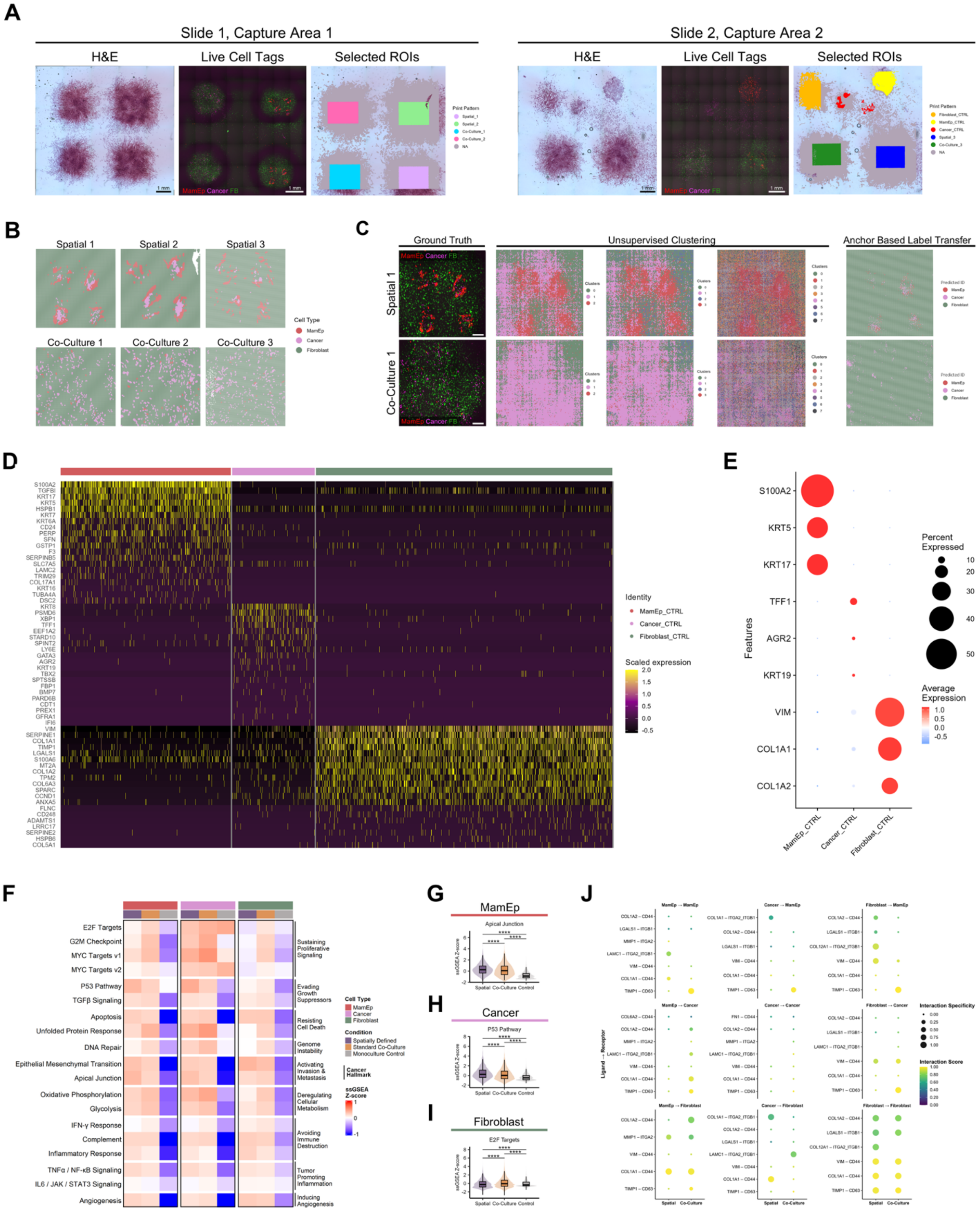


**Figure S7. Spatial transcriptomic analysis of single-cell bioprinted tumor microenvironments spatially defined or randomly deposited co-cultures**

(A) Images of two Visium HD capture areas used for spatial transcriptomic analysis. H&E after 24 hours of culture, live-cell fluorescence image acquired immediately prior to fixation, and selected region of interest used for downstream spatial transcriptomic profiling are shown. Scale bars, 1 mm.

(*legend continued on next page*)

(B) Comparison of barcode annotation approaches. Representative spatially defined and co-culture bioprints shown with corresponding ground truth image; cells were transduced with unique fluorescent reporters prior to printing to enable cell type identification. Scale bars, 200 μm.
(C) Spatial annotation of barcodes for each spatially defined bioprint and co-culture bioprint sample.
(D-E) Differentially expressed genes of each single population control.
(F) Single spot gene set enrichment (ssGSEA) Z-scores for selected MSigDB Hallmark cancer pathways.
(G-I) Distribution of ssGSEA Z-scores for (G) MCF10A, (H) MCF7, and (I) fibroblasts across patterned, random, and single population control conditions. Pairwise differences were assessed by Wilcoxon rank-sum test; * $p < 0.05$, ** $p < 0.01$, *** $p < 0.001$, **** $p < 0.0001$.
(G) MCF10A apical junction ssGSEA Z-scores.
(H) MCF7 P53 pathway ssGSEA Z-scores.
(I) Fibroblast E2F targets ssGSEA Z-scores.
(J) Global overview of ligand-receptor interactions across spatial conditions.

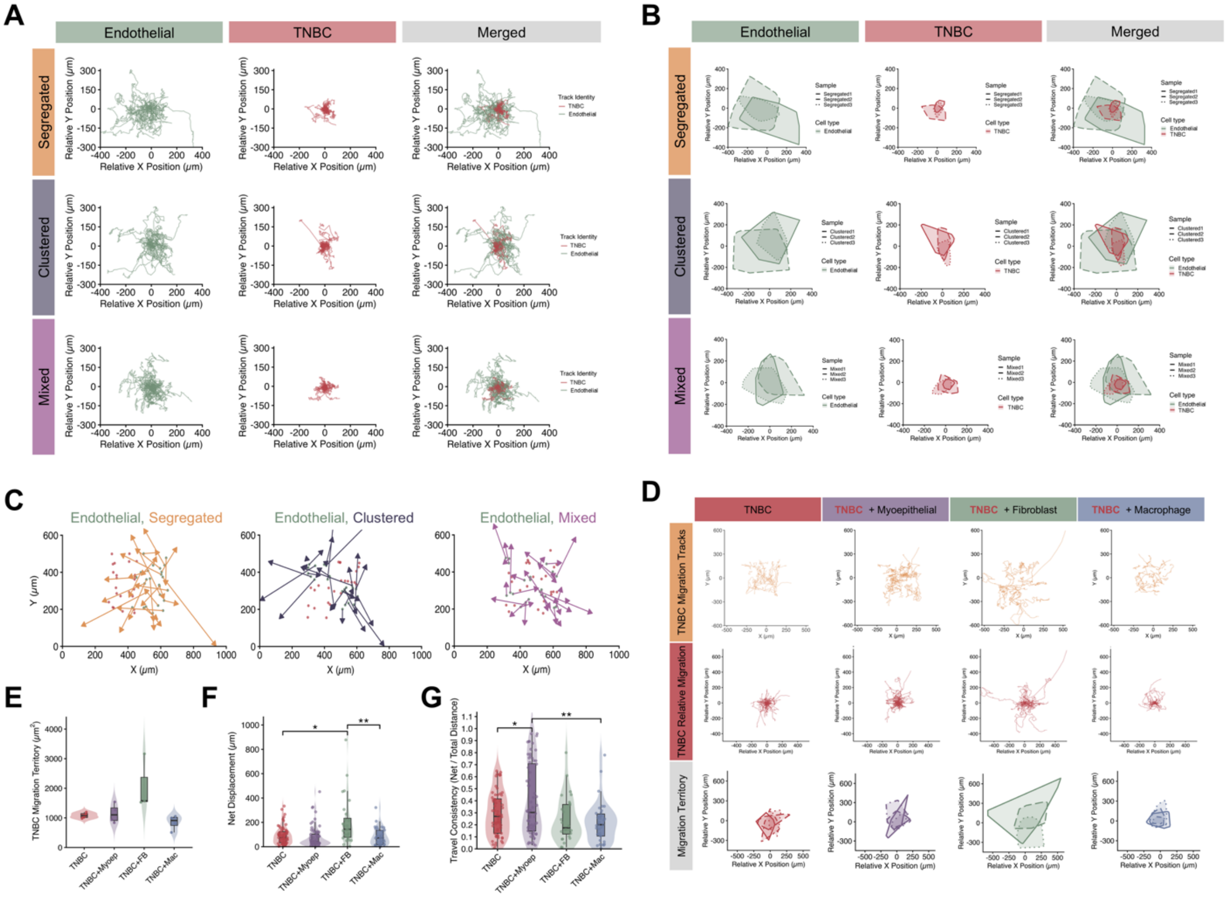


#### Figure S8. Single-cell migration analysis

(A) Relative position migration tracks for HUVEC and TNBC arranged in three bioprint patterns: segregated, clustered, and mixed. $n$ = 3 bioprints.

(B) Migration territory assessment from normalized (relative) position tracks. Bioprint replicates shown, $n = 3$.

(C) HUVEC displacement relative to initial cell positions, replicates merged. Arrow origin denotes starting position, arrow length equals displacement (μm), and arrow point indicates final position. Red spots = MDA-MB-231 and green spots = HUVEC.

(D) MDA-MB-231 migration tracks, relative position migration tracks, and migration territory from relative position tracks. $n = 4$ for all groups except TNBC+FB, $n$ = 3.

(E) Quantification of MDA-MB-231 migration territory.

(F) Net displacement of individual MDA-MB-231 cells. Welch ANOVA with Benjamini–Hochberg corrected Welch pairwise t-tests; * $p < 0.05$, ** $p < 0.01$, *** $p < 0.001$.

(G) Travel consistency (net displacement divided by total path length) of individual MDA-MB-231 cells. Welch ANOVA with Benjamini–Hochberg corrected Welch pairwise t-tests; * $p < 0.05$, ** $p < 0.01$, *** $p < 0.001$.

#### SUPPLEMENT VIDEOS

**Video 1.** Recirculation zone formation under varying internal and external vacuum settings, visualized using fluorescein in a transparent medium.

**Video 2.** Breast cancer biopsy replica with subcellular resolution. Mammary epithelial (red), triple negative breast cancer (pink), mammary fibroblast (no label), mesenchymal stem/stromal cell (no label), and macrophages (no label). Live-cell imaging at 10 minute intervals for 24 hours.

**Video 3.** Spatially defined breast tumor microenvironment model. Mammary epithelial cells (red), triple negative breast cancer (pink), and mammary fibroblasts (no label). Live-cell imaging at 10 minute intervals for 24 hours.

**Video 4.** Randomly bioprinted co-culture model. Mammary epithelial cells (red), triple negative breast cancer (pink), and mammary fibroblasts (no label). Live-cell imaging at 10 minute intervals for 24 hours.

**Video 5.** Cell-cell interaction array using human umbilical vein endothelial cells (HUVEC, GFP+) and triple negative breast cancer cells (red cell tracker) in a segregated spatial arrangement. Live-cell imaging at 10 minute intervals for 30 hours.

**Video 6.** Cell-cell interaction array using human umbilical vein endothelial cells (HUVEC, GFP+) and triple negative breast cancer cells (red cell tracker) in a clustered spatial arrangement. Live-cell imaging at 10 minute intervals for 30 hours.

**Video 7.** Cell-cell interaction array using human umbilical vein endothelial cells (HUVEC, green) and triple negative breast cancer cells (red cell tracker) in a mixed spatial arrangement. Live-cell imaging at 10 minute intervals for 30 hours.

**Video 8**. Cell-cell interaction array of triple negative breast cancer (control). Live-cell imaging at 10 minute intervals for 24 hours.

**Video 9.** Cell-cell interaction array using triple negative breast cancer and myoepithelial cells in a mixed spatial arrangement. Live-cell imaging at 10 minute intervals for 24 hours. Microscope refocused after 1 hour and files combined.

**Video 10.** Cell-cell interaction array using triple negative breast cancer and mammary fibroblasts in a mixed spatial arrangement. Live-cell imaging at 10 minute intervals for 24 hours.

**Video 11.** Cell-cell interaction array using triple negative breast cancer and macrophages in a mixed spatial arrangement. Live-cell imaging at 10 minute intervals for 24 hours.
