## Supplementary Text for "A Single-Cell Bioprinting Method to Reconstruct Native Cellular Microenvironments with Subcellular Resolution"

### SUPPLEMENT TEXT

Our single-cell bioprinting approach uses a microfluidic dispensing system (Biopixlar, Fluicell) that deposits cells, one at a time, suspended in their complete cell culture medium (**Fig. 1B and S1**). The microfluidic device has eight independent wells, four for single-cell suspensions and four for waste collection (**Fig. S1B**). To enable high precision single-cell patterning, the cell must immediately bind to the substrate after being deposited. This is made possible by submerging the printhead in cell culture medium and positioning it <10 μm from the print surface so that as a cell exits the printhead it comes in contact with the underlying substrate. Upon contact, the cell can immediately bind through integrin binding to an extracellular matrix protein and/or electrostatic interaction with a positively charged surface (**Fig. 1B**). If the cell does not immediately attach, it is taken up by external vacuums that flank either side of the delivery channel, along with the cell culture medium that the cell was transported in.

The flow of the microfluidic device is controlled by four independent parameters, the non-delivery pressure (0-50 mbar), delivery pressure (0-450 mbar), internal vacuum (0-300 -mbar), and external vacuum (0-300 -mbar). The non-delivery pressure maintains the flow of cells within all delivery wells where they converge in the crossover zone before interfacing with two internal vacuum lines (**Fig. S1C**). To dispense cells from a desired well, the delivery pressure is applied only to that well to create a no-slip boundary with the other lanes (**Fig. S1D**). The cell-laden medium that passes the internal vacuum is then dispensed out of the printhead where it interacts with the external vacuums. The relationship between the delivery pressure and the external vacuum pressures creates a confined recirculation zone of one miscible liquid inside another (**Fig. 1B and S1D**).

In order to controllably pattern single cells, we first needed to identify the precise balance of flow parameters that would support cell transport and deposition. We began by identifying the conditions that would produce a stable recirculation zone which is required for ideal cell binding during dispensing. We hypothesized that lower delivery pressures would exert less stress on the cells and ultimately lead to greater control over cell placement due to the slower flow speeds. Additionally, we hypothesized the non-delivery pressure would not be necessary and would likely lead to higher incidence of cell crossover (depositing cells from the wrong well). Therefore, we chose to investigate the stability of the recirculation zone at 60, 80, and 100 mbar delivery pressure, with 0 mbar non-delivery pressure, while incrementally varying the internal and external vacuum settings (**Fig. S2**).

The recirculation zone was visualized by continuously dispensing fluorescein into an optically clear medium (**Figs. 1C-D and S2A-C; Video 1**). Recirculation zones that did not expand beyond a defined boundary were deemed stable (**Fig. S2D**). We found that pressure ranges that were able to consistently produce a stable recirculation zone across printheads varied depending on the combinations of delivery pressure, internal and external vacuums. For instance, 80 mbar of delivery pressure formed a stable recirculation zone for an external vacuum of -80 mbar when the internal vacuum ranged from -30 to -40 mbar. When the external vacuum was adjusted to -60 mbar, a higher internal vacuum was needed to offset the reduced vacuum; only -40 mbar internal vacuum was able to form stable recirculation at -60 mbar external vacuum.

We next sought to determine the pressure conditions that would allow cells to be dispensed from the printhead consistently as single cells, as opposed to doublets, triplets, or clusters. Human umbilical vein endothelial cells (HUVECs) were loaded into the printhead and dispensed, while the internal vacuum setting was varied at 5 mbar increments from 0 to -50 mbar, to identify the maximum viable internal vacuum setting (**Fig. 1E and S3A-B**). As predicted, the threshold to enable cell dispensing was less than the threshold to enable fluid dispensing; at 80 mbar delivery pressure and -80 mbar external vacuum, fluid is consistently dispensed up to -40 mbar internal vacuum, but cells are only able to be consistently dispensed up to -25 mbar internal vacuum.

We then characterized the conditions that would enable controlled dispensing of a single cell type when all the dispensing wells of the device were loaded with different cell populations. GFP+ HUVECs were loaded into the desired dispensing well and GFP+ HUVECs that were co-stained with Hoechst were loaded into the remaining wells. If a co-expressing (GFP + Hoechst) HUVEC was deposited, unwanted crossover from another delivery well occurred and we concluded the internal vacuum setting was too low (**Fig. S3C-E**).

The remaining pressure combinations that maintained a stable recirculation zone, enabled cells to be deposited, and prevented unwanted channel crossover were investigated for their ability to controllably pattern single-cells. HUVECs were dispensed onto a collagen substrate and the printability was assessed based on the binding success, external vacuum interaction with underlying substrate, ability to deposit a single cell vs doublet(s), and printing speed (**Fig. S4**). We found HUVECs were able to immediately bind to the underlying collagen upon contact regardless of pressure combination. However, the lowest viable internal vacuum pressure deposited the most doublets and thus did not result in desirable single-cell printing results. Conversely, the highest viable internal vacuum pressures had the most variability across printheads (**Fig. S4A,C**), requiring the most tuning per experiment. High internal vacuum settings also had overall slower bioprint speeds as a greater number of cells are intercepted by the internal vacuum causing less cell to be deposited per second. We also found external vacuum pressures >70 -mbar would aspirate the collagen substrate, blocking the external vacuum and destabilizing the recirculation zone. Other substrates, such as rigid tissue culture plastic with a single layer protein coating, would likely support single-cell bioprints at higher external vacuum pressures.
